# TS2CG as a membrane builder

**DOI:** 10.1101/2025.04.16.649160

**Authors:** Fabian Schuhmann, Jan A. Stevens, Neda Rahmani, Isabell Lindahl, Chelsea M. Brown, Christopher Brasnett, Dimitrios Anastasiou, Adrià Bravo Vidal, Beatrice Geiger, Siewert J. Marrink, Weria Pezeshkian

**Author notes:** Equal Contribution (Fabian Schuhmann, Jan A. Stevens).

## Abstract

Molecular dynamics (MD) simulations excel at capturing biological processes at the molecular scale but rely on a well-defined initial structure. As MD simulations now extend to whole-cell-level modeling, new tools are needed to efficiently build initial structures. Here, we introduce TS2CG version 2, designed to construct coarse-grained membrane structures with any desired shape and lateral organization. This version enables precise placement of lipids and proteins based on curvature preference, facilitating the creation of large, near-equilibrium membranes. Additional features include controlled pore generation and the placement of specific lipids at membrane edges for stabilization. Moreover, a Python interface allows users to extend functionality while maintaining the high performance of the C++ core. To demonstrate its capabilities, we showcase challenging simulations, including a Möbius strip membrane, a vesicle with lipid domain as continental plates (Martini globe), and entire mitochondrial membranes exhibiting lipid heterogeneity due to curvature, along with a comprehensive set of tutorials.

## Introduction

Molecular dynamics (MD) simulations have emerged as a powerful and indispensable tool for exploring biological processes at the molecular scale. Advances in force field development [1–4], simulation engines [5–7], and computing hardware have expanded the scope of MD simulations, from studying individual proteins to complex molecular systems. A frontier direction in the biomolecular modeling field is the simulation of entire cells or cell organelles with molecular resolution, often referred to as in situ simulations [8]. While traditional approaches study isolated biological components, in situ simulations preserve realistic complex molecular interactions similar to their native cellular environment, allowing for synergetic effects to emerge. Achieving such comprehensive simulations might have seemed impossible two decades ago. However, successful simulations of cellular systems, including reduced minimal cells and mitochondrial membranes, among others, have demonstrated the feasibility of this goal [9–13]. Now, that these pioneering achievements have been accomplished, it is time to streamline the process and to broaden the usage of the MD simulations potential. For the simulation of small molecular systems, starting from a random configuration of single lipid-like molecules often suffices to achieve a converged equilibrium state within a feasible simulation time. This is well-established through classic studies, such as lipid bilayer self-assembly within hundreds of nanoseconds [14]. However, constructing initial configurations for cell scale simulations using traditional tools, if even possible, is daunting, time-consuming, and requires many preparation steps. Therefore, a new generation of tools is needed, ones that leverage experimental data and theoretical insights to better initialize simulations so that these can capture the complexity of cellular environments [13, 15–18].

A major bottleneck in modeling large-scale biological systems is the cellular membrane, which exhibits a highly complex organization that emerges over timescales well beyond the feasible timescale of the current MD models. For example, membrane proteins form functional clusters while attracting characteristic lipid shells, and both can separate into distinct domains based on their molecular properties. These organizational patterns are also coupled with the membrane shape, as both proteins and lipids have their own preferred local curvature [19–21]. While MD simulations can potentially capture the formation of these higher-order structures, they require extensive computational resources to reach equilibrium. This becomes particularly challenging in the context of large in-situ models, where multiple such processes would need to occur simultaneously and the required timescale increases nonlinearly with the system size. To study such systems effectively, we must construct initial states that reflect their natural organization as closely as possible by integrating prior knowledge from both experimental and modeling studies. The complexity is also influenced by the membrane shape, which has been discussed in an earlier study [13]. Earlier tools for membrane structure generation primarily focused on circumventing the self-assembly process by pre-assembling lipids into a bilayer configuration. They can generate lipid bilayers in standard shapes, such as flat membranes and vesicles, with a user-defined lipid composition but random lateral organization. Additionally, they often allow the integration of a limited number of proteins [17, 22, 23]. In recent years, a separate class of tools has emerged that can handle arbitrary membrane shapes [24–26]. However, the tools only allow membrane creations according to certain predefined puzzle shapes, do not consider the lateral organization of lipids, require an already generated flat membrane as an input, or cannot place proteins inside or around the membrane. [24–26]. For this purpose, we introduce TS2CG2.0 (**T**riangulated **S**urface to(**2**) **C**oarse **G**rained) as a membrane builder. The program stands out, allowing the design of membrane systems with arbitrary shapes and customizable lipid and protein compositions [13]. Initially, TS2CG was developed for backmapping dynamically triangulated simulation [27, 28] structures into their corresponding coarse-grained (CG) molecular models as an important element for multiscale membrane simulations using, by default, the Martini force field. As TS2CG focuses on the structural placement, the program can straight-forwardly be adapted to handle lipids and structures described in arbitrary forcefields. Furthermore, TS2CG emerged as a robust tool to incorporate experimentally obtained membrane shapes and compositions into the initial configuration of a CG membrane simulation. Yet, a key challenge remains: generating near-equilibrium lateral organization, especially when accounting for the interplay between membrane shape and molecular composition [29–31]. This feature is particularly crucial for modeling the highly curved shapes of organelles, where the membrane shape strongly affects protein and lipid organization and vice versa [9, 29]. TS2CG 2.0 enables precise control over lateral molecular organization during model construction and can be supplied with an arbitrary membrane geometry through triangulated meshes. The new features could enable the generation of more realistic membrane models by incorporating experimental constraints directly into the building process and thus aligning in-silico studies directly with their experimental counterparts.

While we report on case studies using the Martini 3 force field, TS2CG 2.0 has also been successfully used to generate ready to simulate membranes for the Sirah [2] and CHARMM [32] force fields in two specific simple cases (see S6 Non-Martini membranes). However, we leave testing on more complex systems for future studies or interested users.

## Results

### Building Membranes

Building a membrane ready for simulation via TS2CG 2.0 is done in multiple steps, combining different tools and ideas. Simple membranes, however, can now be generated in a single step by utilizing the analytical shape feature of step 3. The program is hosted on GitHub at https://github.com/weria-pezeshkian/TS2CG-v2.0/. Additional documentation can be found at https://weria-pezeshkian.github.io/TS2CG_python_documentation/. The details of the different procedures will be discussed below in the methods section.

#### Step 0 Installation

TS2CG 2.0 is distributed through PyPI, enabling installation via pip. While the software’s computationally intensive operations are implemented in C++, PyPI handles the compilation of these binary components automatically. Users interact with TS2CG 2.0 through Python and command-line interfaces, combining the accessibility of Python with the performance benefits of compiled code.

#### Step 1 Surface Discretization

In the first step, the general shape of the to-be-built membrane is created. For complex membrane shapes, a triangulated mesh is converted into a discrete point representation through the PLM (pointillism) subroutine. The points inherit the attributes of their corresponding surface element, which include a surface area, local coordinate, and curvature tensor. They are generated using a specific algorithm introduced in the first version, which ensures the surface retains its curvature, a critical feature governing membrane behavior at larger scales. PLM accepts triangulated meshes in .tsi format, which can be extracted, for instance, from FreeDTS [28] simulation software output or generated through modeling software like Blender [33–35]. PLM converts the triangulated mesh surface into a discretized set of points and writes a point folder collecting the inner and outer membrane, as well as inclusions (an abstract for proteins location in/on the membrane) and exclusions (an abstract for holes in the membrane). A new extension to PLM is handling shapes that are not closed and allows special treatment of edge cases, see, for instance, the case study Open-edge geometries below. The open edges allow the creation of a different classes of membranes such as nano disks. The points represent surface elements that have spatial coordinates r, area, normal vectors *n*^, and principal curvatures as well as principal directions. The generated point distribution serves as a template for placing molecules during membrane construction while preserving the original surface geometry.

PLM thus transfers a triangle mesh into the point folder, which can be read by the membrane building subroutine (PCG) to construct the membrane structures.

For simple membranes, TS2CG also supports the creation of membranes and direct placement of lipids. A set of analytical geometries are supported, i.e., flat membranes, sine and cosine-shaped membranes, vesicles, or cylinders. This straightforward generation of membranes is implemented in PCG. Additionally, for these geometries, a point directory can be created, which can be used to further modify the membrane composition, as described later.

#### Step 2 Proteins, Pores, and Lipid Domains

TS2CG 2.0 incorporates a Python-based interface for manipulating the generated points. This framework enables precise control over membrane properties through three primary components: membrane composition, Inclusions (protein placement), and Exclusions (membrane pores, which can be introduced to study hole formation and closure). The Point class API allows direct programmatic access for analyzing and modifying membrane composition, enabling informed placement of molecules based on characteristics such as curvature or any user-defined geometric parameter, once the manipulations are handed off to PCG. In case the specific distribution of the inclusions, exclusions, or lipids is not specified, the routine falls back to a random placement. To put the framework into practice, we provide command-line tools that implement common membrane modification strategies, such as creating circular lipid domains around proteins or around any arbitrary point (DAI), assigning lipids based on curvature preference (DOP), or inserting membrane proteins based on curvature preference (INU). These tools enable users to integrate prior knowledge into the membrane composition without the need for Python expertise. The routines are described in detail below.

**DAI:** (“**D**omains **A**round **I**nclusions’) Creates circular domains of a user-defined radius around protein inclusions or arbitrary points. The basic method directly assigns domains based on the Euclidean distance from a user-specified central point. Although this is an efficient implementation, it can produce artifacts if a membrane curves so much, that the different regions of the membrane come closer to each other than the assigned domain radius (see Figure 5 for an example). To resolve this issue, a more computationally intensive, graph-based algorithm is also implemented that employs Dijkstra’s shortest path algorithm [36] to ensure domain continuity within individual membrane surfaces. This latter method prevents artifacts in multi-membrane systems by constructing a distance-weighted network based on the underlying mesh of the distributed points. This tool enables precise control of the membrane composition regarding protein-lipid interactions and the organization of lipid domains. The user chooses which implementation is used when running the program. DAI can assign one domain type per central point but can be run multiple times to add multiple different domains. However, in the case of an overlap to a previous run, the overlapping area will be completely dominated by the last execution of DAI. An application of the procedure is provided in the case study entitled Mitochondrial membrane.

**DOP:** (‘**D**istribution-based **O**ptimized **P**lacement’) Assigns lipid compositions based on the membrane geometry using the local mean curvature. The placement algorithm iterates over each point in the discretized membrane in random order to prevent a systematic bias. For each point, the placement probability for a given lipid type (*l*) is calculated based on the local mean curvature (*H*) and the lipid’s intrinsic curvature preference (*C*_0_) using a Boltzmann-weighted function,

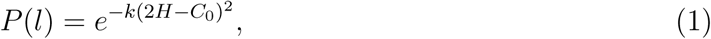

where *k* is a user-defined scaling factor controlling domain sharpness. Additionally, a user can scale *k* further by the area of each point. Through this scaling, the Eq. (1) resembles the Helfrich Hamiltonian. These probabilities, normalized across all lipid types 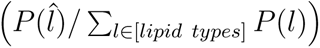, guide the stochastic lipid assignment while preserving the specified overall lipid composition. The curvature-informed placement of lipids allows for the building of membranes closer to equilibrium, substantially reducing the computational cost of the subsequent MD equilibration simulation. The procedure is applied in the case study Integrating Lipid Curvature Preference. It is noted that DOP provides only the weighted function above; however, arbitrary functions are conceivable. In principle, the Point Class API enables complete customization of the probability function used, allowing the implementation of arbitrary lipid sorting schemes tailored to a specific research question.

**INU:** (‘**IN**clusion **U**pdater’) Facilitates easy collision-free protein placement based on local membrane curvature, using the same probabilistic assignment as DOP. For each potential protein position, a placement probability is calculated over the available membrane points using the local mean curvature H, the protein’s preferred curvature *C*_0_, and the user-defined scaling factor *k* that controls specificity. Protein-protein collisions are prevented through a user-defined collision radius. This approach enables the generation of biologically relevant protein distributions across complex membrane geometries. The procedure is used in the case study Mitochondrial membrane.

While these pre-built tools allow useful applications, the underlying Point class API gives users complete programmatic control to develop custom point modification schemes tailored to their specific membrane modeling requirements. Also note, that for most applications, the subroutines through the Python API manipulate the point class and do not generate the membrane. Therefore, execution of the subroutines is significantly faster than the membrane builder described below. Detailed documentation of the Point class implementation is available in Supplementary Material S3 Python documentation.

#### Step 3 Membrane building

The final step in the membrane assembly entails the placement of the lipids and proteins based on the point distribution. The PCG subroutine achieves an overlap-free initial configuration of the membrane system by iteratively placing membrane components according to their spatial requirements, maintaining local geometric relationships and respecting the specified areas per lipid types. A membrane structure (.gro) file accompanied by the respective topology (.top) is returned, ready for simulation with the Gromacs simulation engine [37]. TS2CG, by default, relies on the Martini 3 force field and includes a file (Martini3.LIB) to place most Martini3 lipids. This file can be freely adjusted or replaced to allow placing arbitrary beads, which can then be made meaningful with a custom forcefield which will have to be included in the topology file. A helper program to generate a library entry from a single lipid structure file has been included; *libmaker*. TS2CG 2.0 has been executed successfully with Martini 2 (Martini Globe) and a POPC membrane employing SIRAH beads.

Additionally, given a provided protein structure, PCG can directly place proteins in the membrane. The precise location can be specified through the Point Class API, INU, or a random placement can be achieved. For each protein the z-height can be provided in the configuration input file to have the structure inside the membrane at the right depth or have it hover above or below. With this, it can be adjusted how far a protein might stick out of the membrane or how far it floats above the membrane at the start of the simulation to directly control for physiological conditions. The proteins are placed directly as specified in the input file and no post-placement relaxation is performed. Additionally, a pre-oriented protein structure yields better results as the placement height in z direction is determined from the z direction in the protein structure file. If the specified height is 0, the protein’s center will be in the center of the membrane. Furthermore, if a protein exists in multiple orientations, it can be redefined as multiple proteins in their respective orientation with a separate structure file for each configuration allowing full user control.

Note, that the membranes or membrane protein complexes generated by TS2CG 2.0 serve as an initial structure that ought to be minimized and equilibrated to achieve a realistic membrane (within the bounds of the forcefield).

TS2CG 2.0 also includes practical utilities to simplify the membrane-building workflow. A visualization tool (VIS) enables easy visualization of the point distributions via the command line, allowing validation of domain assignments before membrane construction. Furthermore, the package also provides a solvation tool. This tool works similarly to gmx solvate [37] but is specifically designed for the fast propagation of a small, equilibrated water box into a larger box containing certain particles. It ensures that solvent particles do not overlap with existing particles within a user-defined cutoff radius and also allows for fast placement of ions. A comprehensive tutorial series demonstrating these features and core functionalities is available in the supplementary materials and on the GitHub repository.

### Case Studies

To demonstrate the versatility of TS2CG 2.0, we constructed several membrane systems, some biologically relevant and others that showcase the technical limits of the program’s new features. For each system, we performed MD simulations to validate that the created initial structure is simulation-ready. While the first example highlights the simpler functions of the daily routine of membrane-protein system initialization, the other examples emphasize TS2CG 2.0’s ability to handle arbitrary membrane shapes, complex protein-lipid organizations, and non-standard membrane compositions. An extensive tutorial containing vesicles and flat membranes can be found in the supplementary material (S1 Tutorials).

#### A flat membrane

While the later case studies focus on the more specialized features of TS2CG 2.0, this first case study aims to showcase the creation of a flat membrane with two lipids and a single protein placed slightly above the membrane. For the creation of the presented system, elements from Tutorial 4 and 6 were combined as presented in the supplementary information (S1).

The membrane contains two types of lipids. One type has an occurrence of 90% and an area per lipid of 0.64 *nm*^2^. The other makes up 10% of the lipids with an area per lipid of 0.77 *nm*^2^. The system is amended by a single protein structure, which is placed randomly on the membrane with an elevation of 5 *nm* to the membrane. The thus created initial system was created in 0.87 seconds and is shown in Fig. 2A.

**Figure 1:**
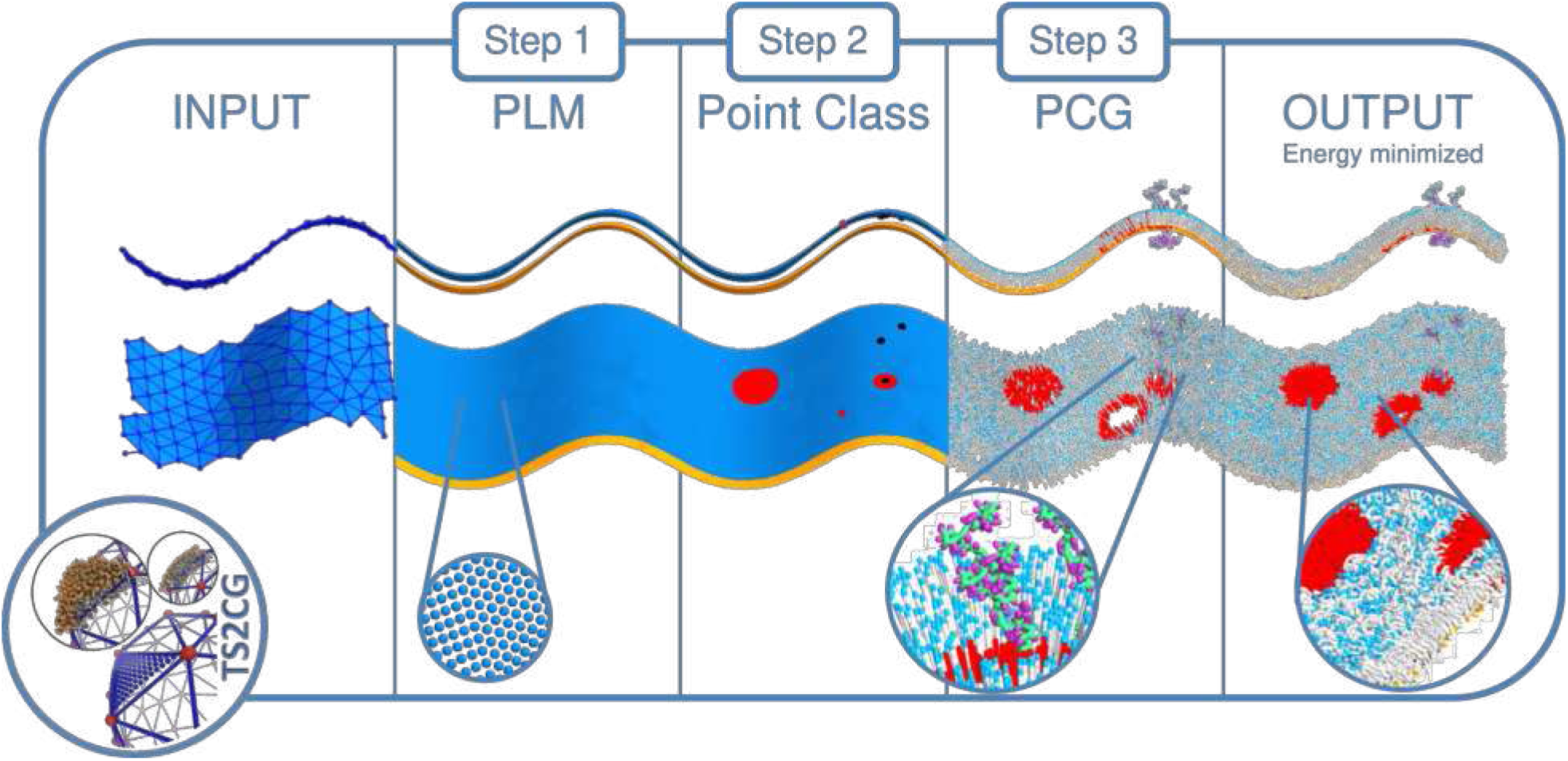
The TS2CG 2.0 membrane creation workflow. A workflow diagram explaining the overall steps of TS2CG 2.0 to generate complex membranes. The process can be started via an analytical shape or an arbitrary triangulated surface. Here, we consider a sin shape. Through PLM or PCG, a point directory can be created which is then manipulated using the Point class to place proteins, exclusions or introduce a different domain (inclusions (proteins) as black dots, domain in orange, and the center of an exclusion is marked in red). A second execution of PCG turns the point folder including the changes into a membrane structure ready for subsequent simulation. PLM and PCG are names for TS2CG 2.0 subroutines, which are explained in the step-by-step guide.

**Figure 2:**
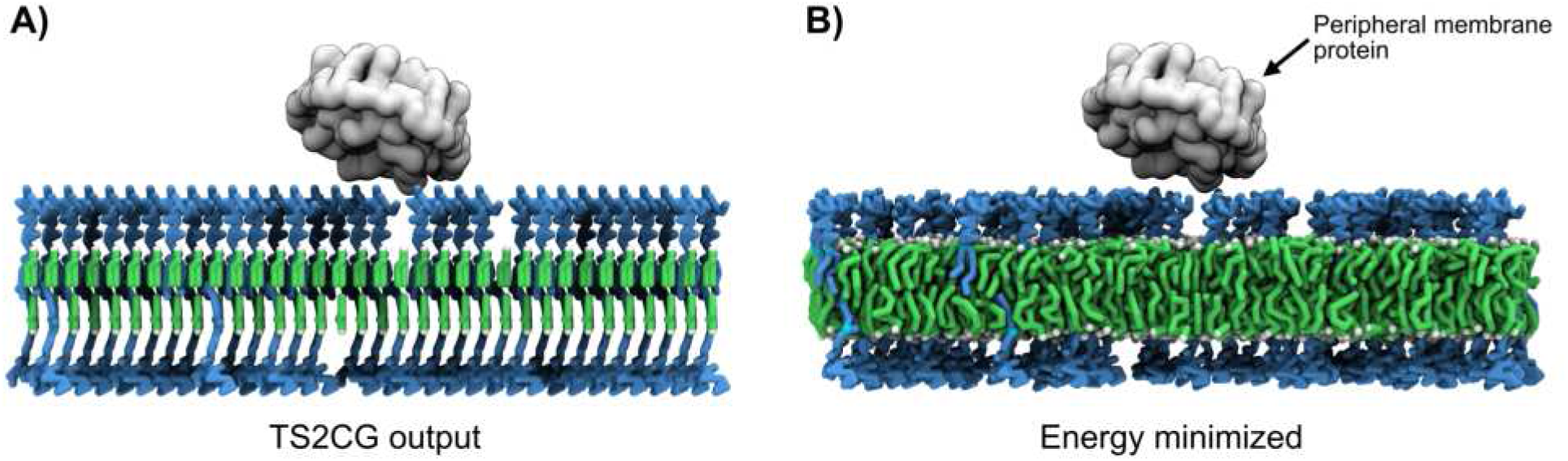
A flat membrane. A membrane with a single protein is shown after initialization through TS2CG (A) followed by a subsequent short MD simulation (B). The image has been centered around the protein.

After a short minimization and equilibration a short simulation was performed akin to the standard simulation setup described in the supplementary methods and the tutorials. The resulting structure (excluding solvent) is visualized Fig. 2B.

#### Integrating Lipid Curvature Preference

Here, different lipids are placed according to the curvature preference profile given in Eq. (1). We show the results for varying TS2CG parameters (*k* and *C*_0_) to place around 20% cardiolipin (CDL) in a curved POPC membrane. The DOP feature to scale *k* further in Eq. (1) is not utilized for this case study. The *C*_0_ value represents the lipid’s curvature preference whereas the *k*-value is a scaling factor controlling domain sharpness. The aim is to generate membranes closer to equilibrium from the start. CDL is known to favor negatively curved regions [29]. Hence, we will work with fixed values of *k* and *C*_0_ for POPC and test how varying *k* and *C*_0_ values impact the placement of CDL. Nine different *C*_0_ values were tested, each with ten different *k*-values, resulting in a total of 90 systems. All test systems underwent a brief energy minimization. We then compare the biased placement of TS2CG with the end result of a 1 *µs* simulation which allowed for the natural sorting of the lipids after an initial random placement of the different lipid types. The simulated self-sorted membrane is considered to be the reference membrane for this case study.

The results are analysed based on two different methods. First by calculating lipid-type community scores to quantify the impact of parameter choices in DOP on the finally generated membrane structure. A second analysis compares the CDL density of a single-parameter set to the self-sorted reference simulation. For further information regarding the methods and scores, we refer to the supplementary methods.

The results from the community scores, are presented in Figure 3 (A), whereas the results from the density distributions comparison are presented in Figure 3 (B). In Figure 3 (A), higher scores indicate stronger internal connectivity within a community. Conversely, lower scores signify more evenly distributed lipids. Hence, the higher the score, the more clear-cut lipid domains are formed on the membrane and the clearer CDL has been sorted to a certain region on the membrane. Three key observations can be made from Figure 3 (A). First, an increase in the parameter *k*, that controls the strength of the bias, results in higher scores. Second, both high and low values of the spontaneous lipid curvature, *C*_0_, contribute to increased scores. As expected, in absence of spontaneous curvature (*C*_0_ = 0), no significant communities are observed, regardless of the *k* value. This is because *C*_0_ is also equal to 0 for POPC, meaning that for every point, both lipid types have the same probability of being assigned, making the assignment effectively random. This only holds true because both lipid types share the same *C*_0_ value. Overall, higher *k*-values result in higher scores and a more segregated membrane. The highest community score is observed for the largest spontaneous curvature tested (*C*_0_ = 0.3), in combination with the largest bias *k* = 10, while another high score is found for *C*_0_ = −0.3, also with *k* = 10. For *C*_0_ = 0.3, PCG places the CDL lipids in the positively curved regions, whereas for *C*_0_ = −0.3, PCG places these lipids in negatively curved regions. The positively curved regions are larger than the negatively curved regions due to asymmetry of the membrane shape. As a result, more lipids can be placed in the positively curved areas, which increases the connectivity within communities, leading to the highest community score. Further analysis will focus on the effect of a higher *k*-value where 20 systems were created with the same *C*_0_ value but different *k*-values. The density data of CDL for each system was extracted and treated as distributions. As shown in Figure 3 (B), the density distributions of these 20 systems were compared to the distribution of the reference system using the Wasserstein distance, where a shorter distance indicates a greater similarity between distributions [38]. The observed trend shows that increasing the k-value leads to a CDL density distribution that more closely resembles that of the reference system. It also shows that there is a limit of how far we bias the placement, the curved areas of the membrane get saturated and PCG cannot place more lipids in the region. The results demonstrate that incorporating DOP into TS2CG 2.0 enables the creation of system conformations closer to a pre-defined equilibrium without requiring long, computationally expensive simulations. While achieving the pre-sorted structures took only minutes on a laptop CPU, the reference structure required hours of equilibration on a high-performance cluster with GPU acceleration.

**Figure 3:**
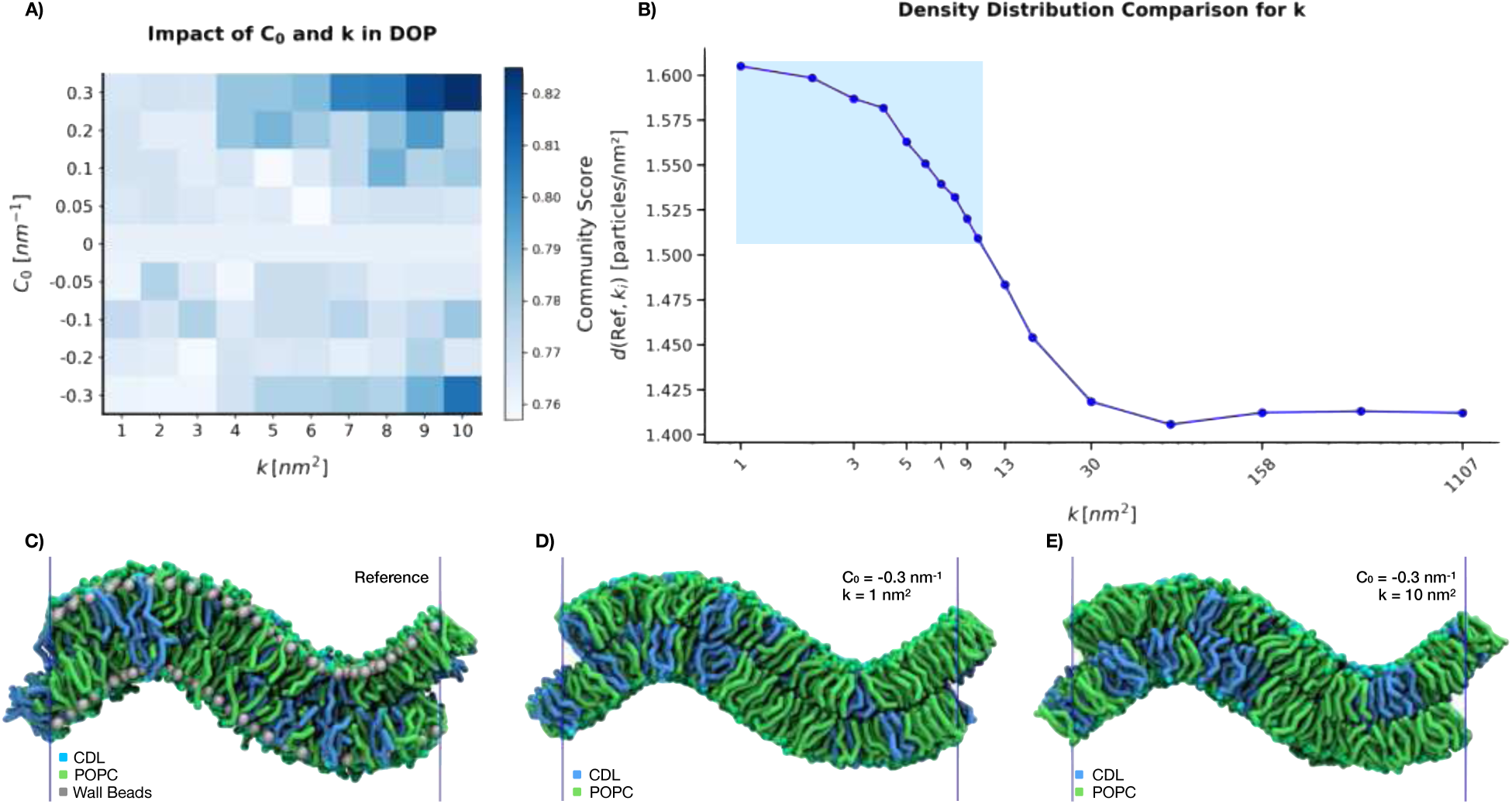
Generating membranes by sorted lipids according to their curvature preference. A) Presents the community scores of the 90 membranes generated with DOP and visualizes the impact of varying *C*_0_ and *k* values for CDL while *C*_0_ and *k* remain constant for POPC (*C*_0_ = 0*, k* = 1). B) Density distribution comparison for different *k*-values, twenty systems generated with *C*_0_ = −0.3 and different *k*-values are evaluated against the reference. The blue box highlights the *k*-values used for the community score. C) The reference system, with natural sorting of the lipids through simulations. D) Resulting system for *C*_0_ of −0.3 and *k* of 1. E) Resulting system for *C*_0_ of −0.3 and *k* of 10. The differences between Panels D and E become best visible in the right valley. The high *k* placement of Panel E places a collection of CDL precisely, while the low *k* placement does not. The blue lines in Panels C to E resemble the periodic boundary condition.

#### Mitochondrion

To explore the integrative modeling strengths of TS2CG 2.0, we built two complementary mitochondrial models: a detailed crista junction based on earlier work by Brown et al. [9] with experimentally informed protein and lipid distributions and a large-scale inner mitochondrial membrane. The first showcases how experimental and simulation data can be integrated into a complex membrane model, while the second presents the software’s capacity to handle mesoscale membrane geometries.

Membrane curvature plays a crucial role in organizing both lipids and membrane proteins within the mitochondrial inner membrane [39], maintaining its complex membrane geometry. In both models, lipids are placed according to experimentally and computationally determined curvature preferences [29]. Specifically, POPC and POPS were preferentially placed in areas of positive curvature (*C*_0_ = 1 *nm*^−1^, *k* = 250 *nm*^2^), while SAPE and PAPI were placed in areas of low curvature (*C*_0_ = 0 *nm*^−1^, *k* = 250 *nm*^2^), and CDL2 placement was biased towards negatively curved regions (*C*_0_ = −1 *nm*^−1^, *k* = 250 *nm*^2^). The realized lipid distribution in the model of the inner mitochondrial membrane, built using an experimentally obtained membrane geometry [40], shows a strong correlation with the curvature of the input structure (Figure 5A,B). Analysis of the relative lipid distributions shows the expected enrichment patterns in accordance with each lipid’s curvature preferences (Figure 4C). The generation process of the whole inner mitochondrial membrane took roughly ten minutes with TS2CG 2.0.

**Figure 4:**
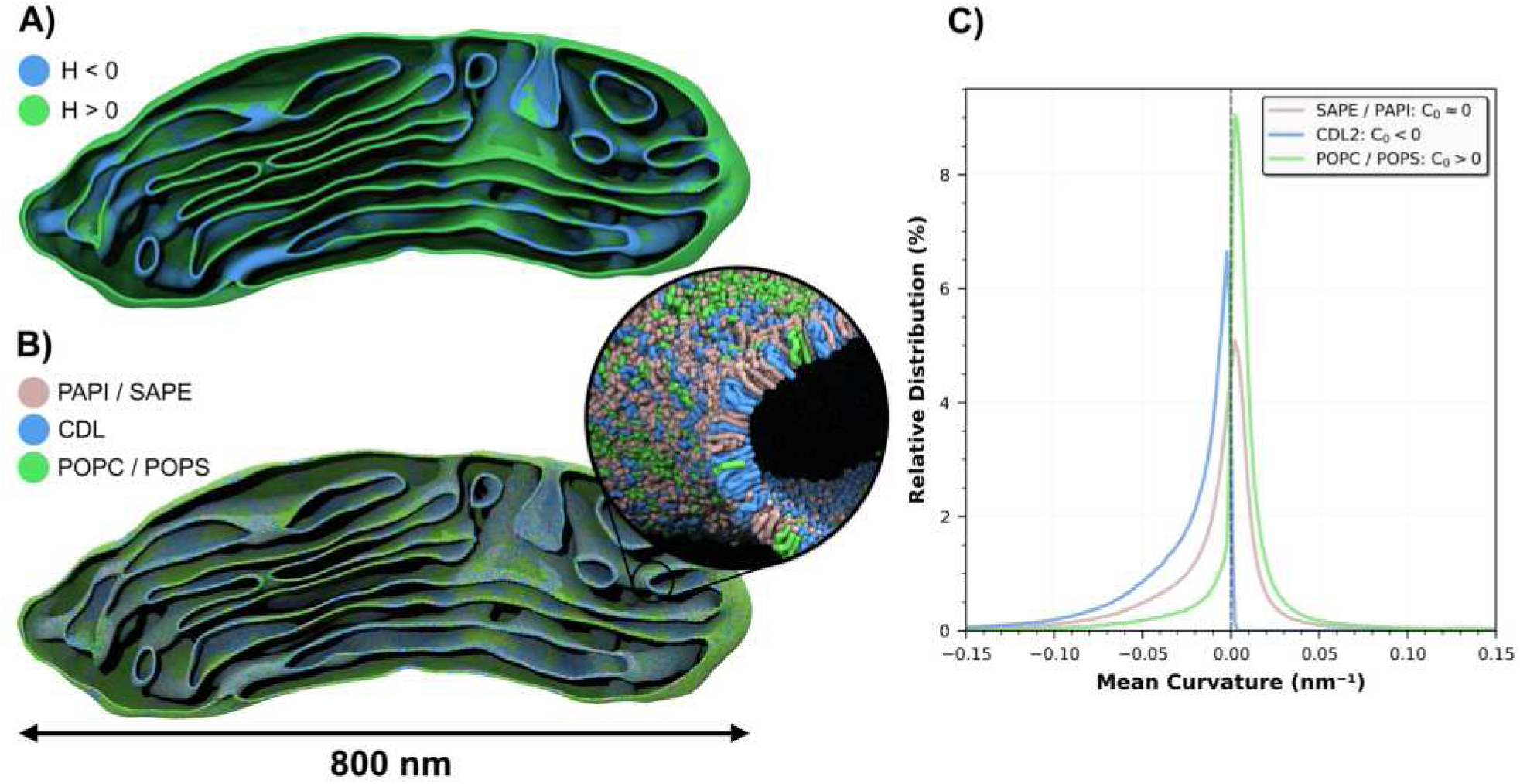
Mitochondrial membrane with real shape from Cryo-ET. A) Inner mitochondrial membrane colored by mean curvature: positive mean curvature (*H >* 0) in blue, negative mean curvature (*H <* 0) in green. B) Martini 3 model of the inner mitochondrial membrane with lipids distributed by curvature preference (PAPI/SAPE pink, CDL blue, POPC/POPS green). C) Curvature-dependent distribution of mitochondrial lipids plotted as the relative distribution in function of membrane curvature. A large *k* was chosen to strongly bias the lipids for a clearer visualization of the tool. Therefore, the curves appear skewed as it is highly unlikely for a lipid to be placed in the region opposite its own preference.

**Figure 5:**
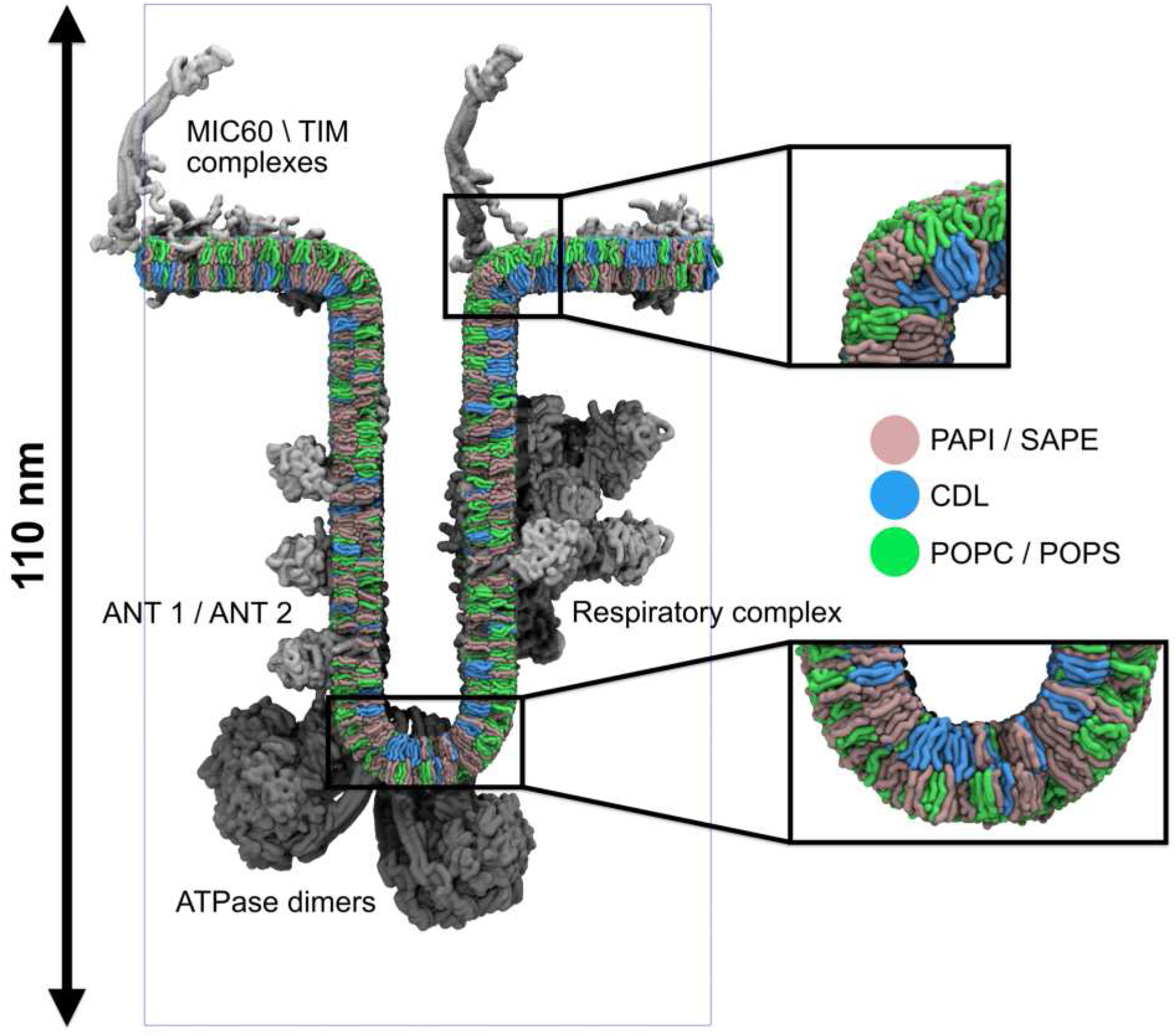
Model of a mitochondrial cristae. The model shows lipids and membrane proteins arranged by curvature preference. MIC60/TIM complexes (top), ANT1/ANT2 transporters and respiratory complex (sides), and ATPase dimers (bottom) are shown. Zoom panels highlight the lipid sorting patterns: PAPI/SAPE (pink), CDL (purple), POPC/POPS (green).

The mitochondrial membrane proteins are similarly organized based on curvature preference. In order to create a relevant in situ model of a mitochondrial crista, we position eight different types of inner protein complexes according to experimentally determined localization on a triangulated surface representing a crista junction [9]. The placement of each protein complex is biased based on specific restrictions on both local mean curvature and z-position to represent its physiologically relevant localization.

As ATPase dimers are found at the crista ridge [41], these complexes were placed on vertices with negative curvature exclusively. The respiratory mega-complexes, ANT1 and ANT2 are found on the crista sides [42, 43] and hence were placed on flat regions of the membrane below the crista junction. The MIC60, TIM22, and TIM23 complexes are found on the inner membrane facing the outer mitochondrial membrane [44, 45], and hence these complexes were also placed on flat membrane regions but with a higher z-position restriction. The numbers of each protein included reflect concentrations reported in the literature [46].

The result of this procedure (Figure 5) is an accurate representation of a complex membrane system, showing curvature-based organization of both protein complexes and lipids based on curvature and membrane position. This enables the construction of complex systems with components placed in such a way that reflects those found in vivo, reducing the amount of simulation time needed to obtain meaningful results.

#### Open-edge Geometries

TS2CG 2.0 introduces support for open-edge meshes, further expanding its scope of application. Supporting open edges enables the construction of both biologically relevant systems (e.g. nanodiscs) and abstract geometries that were previously difficult or impossible to build.

To illustrate the power of this capability, we present a Möbius strip lipid membrane. The Möbius strip presents a challenging geometry as a non-orientable surface with a single continuous edge. The non-orientable nature results in a discontinuity in the surface normals across the mesh. As such an input mesh will always be cut through to recreate the orientable surface, and thus leading to open edges in all directions. TS2CG 2.0 can now incorporate this open-edge geometry and produce a Möbius strip membrane.

We constructed two versions of this Möbius membrane to showcase the edge-based lipid placement available in the new Python API (Figure 6). The first membrane consists of a simple binary mixture of 30% cholesterol and 70% POPC lipids, randomly placed. When simulated, the membrane’s line tension drives a reduction in edge length, with the system evolving from the initial Möbius strip geometry to an intermediate Sudanese Möbius configuration (where the edge forms a circle, minimizing line tension energy) and finally to an oblate vesicle over a few hundred nanoseconds of CG simulation time. For the second version, we identified edge vertices in our mesh and selectively placed specific lipids at these locations. Since shorter lipids might stabilize the high curvature at the membrane edge, we placed DLPC lipids specifically along the edge vertices. The two systems showed different kinetics, although both eventually formed oblate vesicles. While the POPC/CHOL system completed its transition by 300 ns, the addition of DLPC noticeably delayed this process by a 100ns. Of course, to validate the statement statistical replicas would need to be conducted, which is beyond the scope of this proof-of-concept. For quantitative analysis of the topological transitions using leaflet segmentation, see Supplementary Figure S1. These simulations serve primarily as proof of concept for the technical capabilities of TS2CG 2.0, illustrating how programmatic edge-specific lipid placement can be applied.

**Figure 6:**
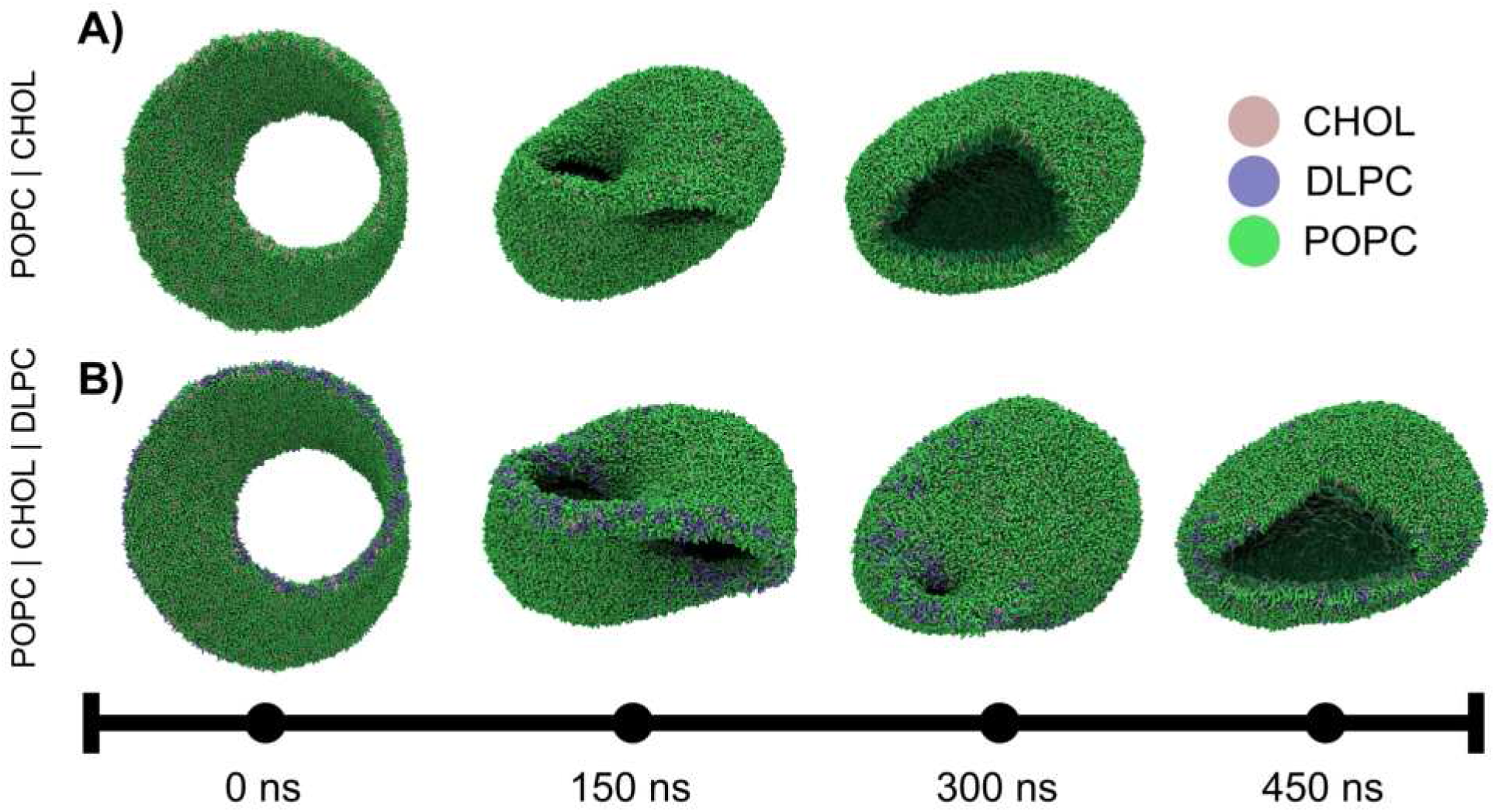
The Martini Möbius strip. Representative snapshots of lipid Möbius strips transforming into vesicles. A) 70%POPC /30%CHOL composition transitioning to an oblate vesicle with complete bilayer formation within 300 ns. B) 68%POPC /29% CHOL/3% DLPC composition, where DLPC was placed in the open edges resulting in a delayed bilayer formation occurring at 400 ns. Cutouts reveal the topological transition from a single continuous monolayer to two separate lipid bilayer leaflets in each oblate vesicle.

Another application of open-edge geometries results from the periodic boundary conditions used in MD simulations. Membranes that form continuous structures in periodic boundary conditions appear as discontinuous, open-edged structures in Euclidean space. Since 3D modeling software typically operates in Euclidean space, meshes that should be continuous when simulated with periodic boundaries are frequently exported as open structures. Our new implementation addresses this limitation by properly handling these open-edged geometries within the periodic simulation environment. Lipid cubic phases are examples of such systems, where a membrane lying on a triply periodic minimal surface divides space into two bicontinuous water channels [47, 48]. Self-assembly MD simulations of single-unit cells of lipid cubic phases have provided insights into their structure and the dynamics of their components [49–51]. However, building larger systems composed of multiple-unit cells has remained challenging [52, 53].

The triangulated mesh representing the diamond cubic phase was generated using the analytical solution of the Schwarz diamond minimal surface [54] employing the code supplied by Brasnett et al. [55]. Using monoolein at a water/lipid ratio of 0.29 w/w and 300 K, we constructed two systems, a single unit cell (±10 *nm*^3^) and a 2 × 2 × 2 periodic array of unit cells (visualized in Figure 7). Each system was simulated for 2 *µs*, throughout which the bilayer maintains close alignment with the mathematical minimal surface (Supplementary Figure SF2). The connectivity of the water channels was also analyzed using containment analysis, which showed that the non-intersecting water channels were also preserved (Supplementary Figure SF3) [56]. This model shows that TS2CG 2.0 enables the modeling of increasingly complex membranes, opening new avenues for studying systems from protein crystallization matrices to organelle membrane networks, where cubic phases play crucial biological roles [57–62].

**Figure 7:**
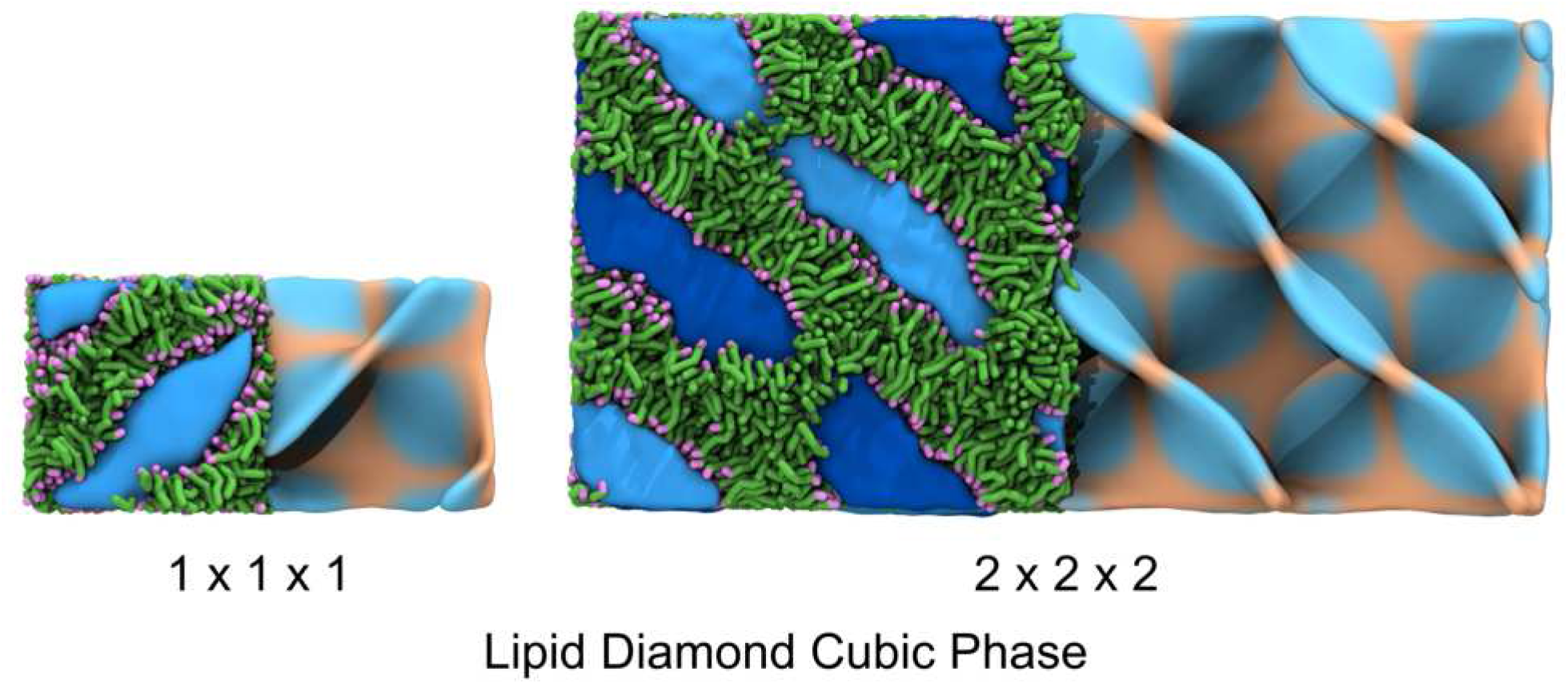
Monoolein diamonds. Snapshots of a monoolein diamond cubic phase at two different lattice sizes (1 × 1 × 1 and 2 × 2 × 2). The left panels show the molecular representation with lipid tails in green, headgroups in purple, and the water channels displayed in hues of blue. The right panels display the fitted surface of the cubic phase lipids colored according to their Gaussian curvature.

#### Martini Globe

In the last case study, the precision of the lipid placement is shown on an artificial case, in which the continents of our planet are mapped onto a membrane vesicle. In the final case study, although it lacks biological relevance, the precision of lipid placement is exemplified using an artificial example where Earth’s continents are mapped onto a membrane vesicle. We used PLM to create a discrete point distribution from a spherical surface mesh. Each point’s coordinates are converted to longitude and latitude, which are then mapped to continental boundaries using public geographic data obtained from US Geological Survey [63] and read by the regionmask python package [36, 64]. Points are assigned to a continent, i.e. lipid domain, based on their geographic coordinates, with oceans designated as domain 0. To distinguish the continental regions, we assign different lipid types to each domain. To demonstrate the fine-grained control of our method, we chose the system size so that Denmark, a relatively small country, can be represented by its own lipid domain. In the final model constructed, Denmark proper is defined by exactly three lipids. The Kingdom of Denmark is then assigned its own domain.

After the globe was created using TS2CG 2.0, it was equilibrated using dry martini [65]. The resulting model is shown in Figure 8. The details for the simulations can be found in the supplementary methods (S4 Methods). An artistic movie representation of the globe can be found in the supplementary material (S5).

**Figure 8:**
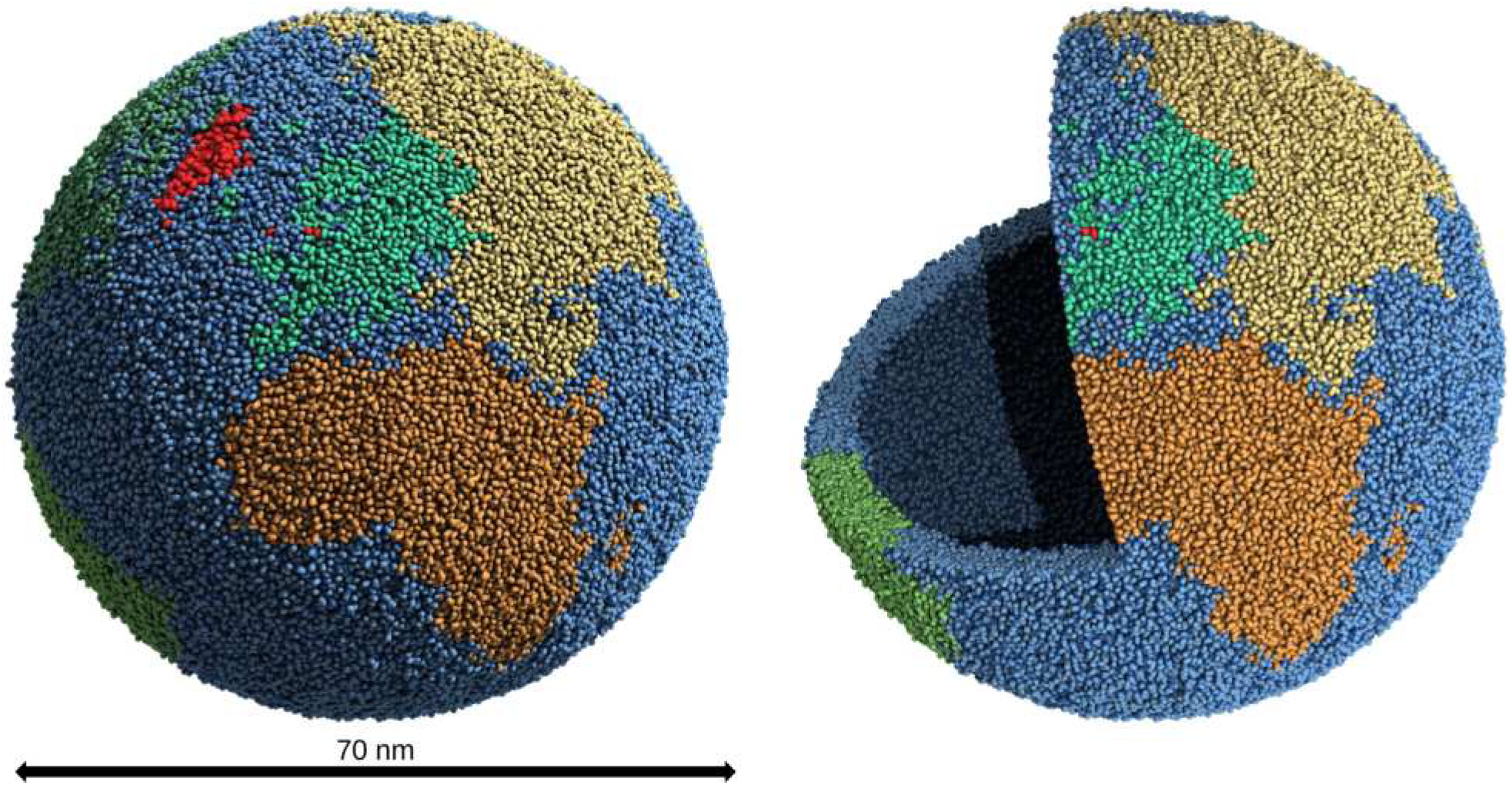
The Martini Globe. The lipids of the globe colored by their continent based on the tectonic plates. The Kingdom of Denmark is colored dark red. Panel A shows the whole surface. Panel B shows a cutout of the globe visualizing the inner and outer monolayer and the lipid tails in between.

### Discussion and Conclusion

TS2CG, version 2.0, introduces significant extensions to its predecessor, emerging as a versatile coarse-grained membrane builder. This new version represents a major advance in membrane system preparation for molecular dynamics simulations. During the development process, alpha-versions of TS2CG 2.0 were already successfully used in other projects to simulate lipid nanoparticles [66, 67].

TS2CG 2.0 now supports open-edge geometries, removing the previous limitation to closed surfaces. This advancement enables easier compatibility with mesh generation tools like Blender, GAMer, and Geogram [33, 35, 68], expanding the range of feasible membrane architectures. We demonstrated the software’s versatility through applications ranging from complex biological structures like mitochondrial membranes to specialized geometries such as lipid diamond cubic phases and a Möbius strip.

For simpler membrane architectures, TS2CG 2.0 introduces analytical membrane generation, enabling easy construction for a set of simple membrane shapes without requiring input meshes. These analytically generated membranes can be coupled with position restraints that preserve their precise geometry throughout MD simulation, which are referred to as wall beads (visualized in Figure 3C). This membrane scaffolding approach enables the systematic investigation of curvature-dependent lipid and protein sorting, generating new insights that inform the construction of more complex membrane architectures.

At its core, TS2CG 2.0 enables precise control over membrane lateral organization through both direct manipulation and curvature-based probabilistic approaches. The direct control over lipid organization is shown through the construction of a model vesicle that precisely maps different lipid types to Earth’s continents, achieving a fine enough resolution to define Denmark proper with just three lipids. While this globe model serves as a technical demonstration, the same methodology enables the construction of biologically relevant membrane organizations, particularly generating phase-separated domains with geometric control. Note that the CG membrane models built by TS2CG can subsequently serve as templates for all-atom simulations through established backmapping protocols [69–71], also providing a robust pathway for generating complex membrane architectures at atomic resolution.

The biological relevance of these capabilities is exemplified in our mitochondrial membrane models. For the large-scale mitochondrial membrane, we showcase the curvature-based lipid distributions, while in the smaller cristae model, both membrane proteins and lipids are positioned according to their specific curvature preferences. This geometry-aware approach to membrane building enables the generation of closer-to-equilibrium starting configurations, increasing the stability of large-scale systems while reducing equilibration times in molecular dynamics simulations.

While TS2CG 2.0 provides control over membrane construction, its capabilities are ultimately limited by available molecular information. Our building protocol requires specific inputs about curvature preferences and molecular distributions, which are often only partially available from experiments. However, this limitation highlights an opportunity in integrative modeling. By combining TS2CG with experimental data from multiple sources - global proteomics, lipidomics, and structural studies - we can explore molecular-scale membrane organization that satisfies these experimental constraints. This approach provides insights into membrane architecture beyond the resolution of current experimental techniques. Future developments will likely incorporate machine learning approaches to predict membrane organization, trained on existing simulations of simpler systems, further bridging the gap between global cellular measurements and molecular-scale architecture. When combined with molecular dynamics simulations, this integration would effectively create a true computational microscope for investigating membrane structure and dynamics in situ [8].

Performance is always a key concern for the computational modeling community. Here, we brought the performance of TS2CG 2.0 to light by constructing the entire inner mitochondrial membrane using curvature-biased lipid placement. Despite the current version of TS2CG being programmed to utilize only a single thread, it can still generate complex membranes within a feasible time frame. For instance, the mitochondrial membrane was created within 10 minutes. However, several features of the PLM and PCG routines could be optimized for multi-threading, enabling even larger membrane-building projects in the future. Implementing these improvements will require a refactoring of the C++ code base, which is already planned in our continuous development pipeline.

In conclusion, TS2CG 2.0 provides a user-friendly workflow that allows users to generate membranes of any composition to initialize complex coarse-grained membrane simulations. In short, TS2CG 2.0 makes constructing intricate membranes possible while also simplifying the creation of commonly used flat membrane simulations.

## Supporting information

Details to S1

Details to S2

Details to S3

## Acknowledgements

This research is supported by the Novo Nordisk Foundation (grant No. NNF18SA0035142 and NNF22OC0079182, NNF20OC0063808). W.P. is also supported by Independent Research Fund Denmark (grant No. 10.46540/2064-00032B) and acknowledges the funding from Marie Skodowska-Curie Fellowship (grant No. 101104867). S.M. and C.M.B. acknowledge funding from the ERC with the Advanced grant 101053661 (“COMP-O-CELL”). S.M. and J.A.S acknowledge funding from NWO through the NWA grant “The limits to growth: The challenge to dissipate energy”.

## Supporting Information

- Main supplementary document: Includes an overview of all available supplements (S1 - S6) and directly includes the methods and supplementary case study information.
- S1: A full-length pdf version of the TS2CG 2.0 tutorials
- S2: The general manual for TS2CG 2.0
- S3: The documentation for the newly developed python functions
- S5: An outreach video of the martini globe simulation (played in reverse)

---

**Figure 9:**
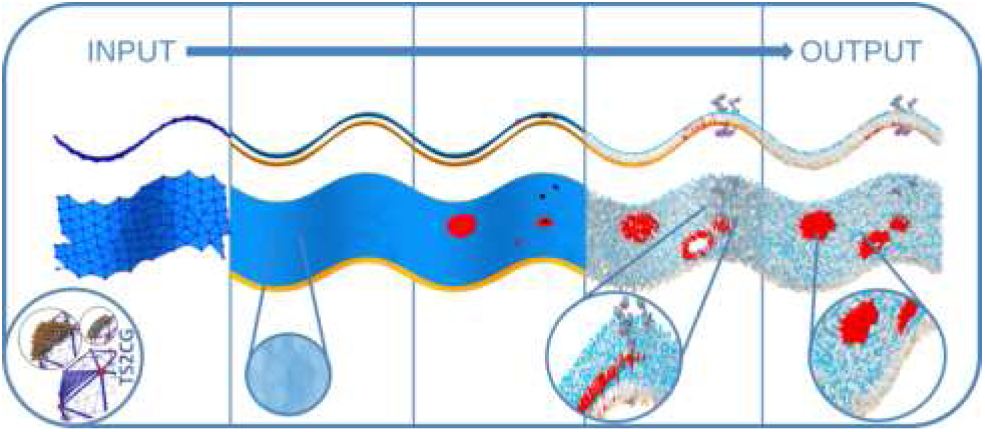
For Table of Contents Only

