## Supplementary Information S1-S6 for "TS2CG as a membrane builder"

### TS2CG as a membrane builder – Supplementary Information

#### S1 Tutorials

See separate supplementary material. S1_Tutorials.pdf

#### S2 Manual

See separate supplementary material. S2_Manual.pdf

#### S3 Python documentation

See separate supplementary material. S3_Python_documentation.pdf

#### S4 Methods

##### MDs settings and analysis

All membrane models were constructed with TS2CG v2.0, and the simulations performed in this paper were run with GROMACS 2023.3 [6] with the Martini 3 force field[1,29].
The MD simulation used the leap-frog integration algorithm and the Verlet cut-off scheme. For nonbonded interactions, the short-ranged Lennard-Jones interactions were truncated at 1.1nm, while long-range Coulomb interactions were treated using the reaction-field method with the same 1.1 nm cut-off. Temperature was maintained at 310K using the velocity-rescale thermostat. For constant pressure simulations, the Berendsen barostat was used during equilibration, while the

c-rescale or Parrinello-Rahman barostat was employed for production runs. The pressure was maintained at $1\ bar$ using a time constant of 12ps and a compressibility of $3\times 10^{-4}\ bar^{-1}$.

Unless otherwise specified in the case study methods sections below, we used a standard Martini3 simulation protocol consisting of the following steps. Initially, the systems were energy-minimized using the steepest descent algorithm. When necessary, double-precision calculations or soft-core potentials were utilized during energy minimization to resolve high-energy contacts. Following the minimization, the systems were solvated using TS2CG's solvation (SOL) subroutine with physiological salt concentration (150 mM NaCl). The solvated models were equilibrated in the NPT ensemble using a 10fs time-step. Subsequent production simulations were performed with a standard 20 fs time-step.

Analysis of the simulation trajectories was performed using Python3 with the MDAnalysis (version 2.7.0)[72,73] and Networkx (version 3.1)[36] packages. Visualizations of the simulation snapshots were created using VMD (version `1.9.4a57`)[74], and the figures were compiled with GIMP (version 2.10.38) and Matplotlib (version 3.7.5)[75].

#### Case Studies

##### Integrating Lipid Curvature Preference

First, the reference system was generated and simulated. The system is a curved POPC lipid bilayer with 20% CDL using TS2CG PCG (version 2.0). The new function DOP is not used when generating the reference structure, resulting in the lipids being placed randomly throughout the bilayer. Wall beads were used to maintain the curved shape of the membrane during the simulation. The wall beads are placed at the level of lipid head groups and interact exclusively with the lipid acyl chain beads in a repulsive manner[29]. The wall beads were kept frozen during the simulation using freeze-groups. The molecular dynamics simulation runs for 1 µs.

The test membranes were created using the DOP method; a total of 90 bilayers with the same shape and lipid composition as the reference structure were generated with varying $C_0$ and k-values for CDL. The $C_0$ for POPC is set to 0.0 for all simulations, while the $C_0$ values for CDL were set to 0.3, 0.2, 0.1, 0.05, 0, -0.05, -0.1, -0.2, and -0.3, respectively, and each was tested with a set of k-values ranging from 1 to 10 in integer steps. In order to make results more comparable, a random seed was set for all membranes. After the membranes are constructed, they are only energy minimized for 20,000 steps.

The first analysis focuses on calculating a community score for each of the 90 membranes. Initially, a pairwise distance matrix is generated based on the coordinates of the headgroups of both lipid types represented by their PO4 beads. The two different lipid types are considered to be two communities, respectively, and they are created using MDAnalysis. A dynamic distance threshold is put into place to generate a network of the PO4 beads such that the network contains exactly two connected components, which then divides the membrane into its upper and lower monolayer. The analysis is then performed for each monolayer separately. Separated distance matrices per monolayer are calculated from which graphs are constructed where nodes represent PO4 beads (the positions of the two PO4 beads in CDL are averaged to create one location per lipid) and edges are set according to a distance threshold of 15 nm with the distance itself serving as a weight.

The two communities, POPC and CDL, are analysed within each monolayer. To assess the quality of the partition in the distance graph, the partition function “partition_quality” in NetworkX is used, evaluating the distribution of intra-community and inter-community edges[76]. This function returns the *coverage*, which quantifies the proportion of edges contained within the communities relative to the total number of edges, serving as the score. The score for each monolayer was averaged, and the results can be seen in Fig. 3(A).

The second analysis focuses on the impact of varying k-values. A total of twenty systems were created using a $C_0$ value of -0.3 $nm^{-1}$ and a set of k-values: 1, 2, 3, 4, 5, 6, 7, 8, 9, 10, as well as $10+e^i$ for i=1,2,…,10. Density maps for CDL were generated for the twenty bilayers, as well as the reference membrane, using GROMACS's built-in function “gmx densmap”. The resulting density maps were unified to a common height and width, and the density data were treated as distributions of CDL. Then, the Wasserstein distance was applied through the Python function “wasserstein_distance” to compare the distributions obtained from the membranes, evaluating how the distributions from membranes generated with different k-values differ from the reference distribution. In Fig. 3(B), the calculation is denoted as *d*(Ref, k_i), where *d* represents the Wasserstein distance, 'Ref' refers to the density distribution from the reference, and k_i corresponds to the distributions obtained from membranes created with different k-values. Between distributions obtained from the membranes. The Wasserstein distance, also known as Earth Mover's Distance, measures the difference between two probability distributions by quantifying the minimal cost required to transform one distribution into the other. In this context, it provides a measure of how closely the CDL density distribution of each system aligns with the reference distribution, with smaller distances indicating greater similarity. The results are shown in Fig. 3(B).

##### Mitochondrion

Two distinct mitochondrial membrane models were constructed: a large-scale lipid-only representation of the inner mitochondrial membrane and a more detailed cristae junction with protein complexes.

For the inner mitochondrial membrane, we utilized experimentally obtained mesh from Pezeshkian et al.[13] The lipid composition incorporated key mitochondrial lipids in the ratio POPC:SAPE:PAPI:POPS:CDL2; 29:36:5:3:26. We employed curvature-based lipid placement, assigning CDL2 a negative curvature preference, while POPC and POPS were given positive curvature preferences. To create a strong bias for the lipids to sort to their preferred curvatures, we used a scaling factor (k) of 250. Using PCG, only the inner leaflet of the membrane was created to better highlight the correlation between lipid placement and the curvature map of the input mesh. The resulting model was equilibrated through a sequence of soft-core and double-precision energy minimization, followed by short 5ns implicit solvent equilibration using the stochastic dynamics integrator using a timestep of 5fs. The energy minimization and brief equilibration in vacuum serve to allow the lipids generated by TS2CG to adopt their native conformational states. This is necessary for creating geometrically valid lipid structures before visualizing the system.Only one leaflet was built since visualizing both leaflets is not helpful for highlighting regions of high and low curvature. The result is clearer when only one leaflet is shown.

The cristae junction model was based on a triangulated mesh from previous work by Brown et al.[9].The mesh was converted to a discretized point cloud using the Pointillism subroutine with a bilayer thickness of 3 nm. We applied the same curvature-based lipid placement strategy but here we used a scaling factor (k) of 100. Protein complexes were placed in the crista model by selecting inclusion points based on their known curvature preferences and spatial localization in mitochondria. The z-dimension was used to distinguish proteins on the crista sides from those at the crista ridge. ATP synthase dimers were placed exclusively at regions of negative curvature and lower z-position, corresponding to the crista ridge, while respiratory complexes were positioned along the flat regions of the crista sides. Membrane proteins associated with the inner boundary membrane were restricted to flat membrane regions with higher z-position values. An exclusion radius was defined for each protein complex to prevent steric clashes. The final membrane structure was constructed using PCG with a bond length parameter of 0.2 nm. The system was briefly equilibrated using the standard Martini 3 simulation protocol described above.

##### Open-edge Geometries

The Möbius strip lipid membrane was constructed using a triangulated mesh modeled in Blender (version 4.2). The mesh was processed using the Pointillism subroutine to generate a discretized point cloud representation with a bilayer thickness of 2 nm.

Two distinct versions of the Möbius strip membrane were created to highlight our new edge attribute in the TS2CG Python API. The first version consisted of a simple bilipid composition of 30% cholesterol (CHOL) and 70% POPC. For the second version, we aimed to stabilize the exposed membrane edge by precisely placing DLPC lipids along the edge vertices of the mesh, the resulting relative concentrations are 29% cholesterol, 68% POPC, and 3% DLPC.

Both membrane systems were simulated following the standard protocol described above, with production runs extending to 500 ns. Throughout the simulations, we tracked the number of lipid leaflets to distinguish the Möbius and vesicle state of the membrane[77]. Although we didn't succeed in fully stabilizing the Möbius membrane, the addition of DLPC lipids at the membrane edge significantly delayed the topological transition compared to the simple bilipid mixture. It should be noted that these simulations were performed as single replicates and serve primarily as proof of concept.

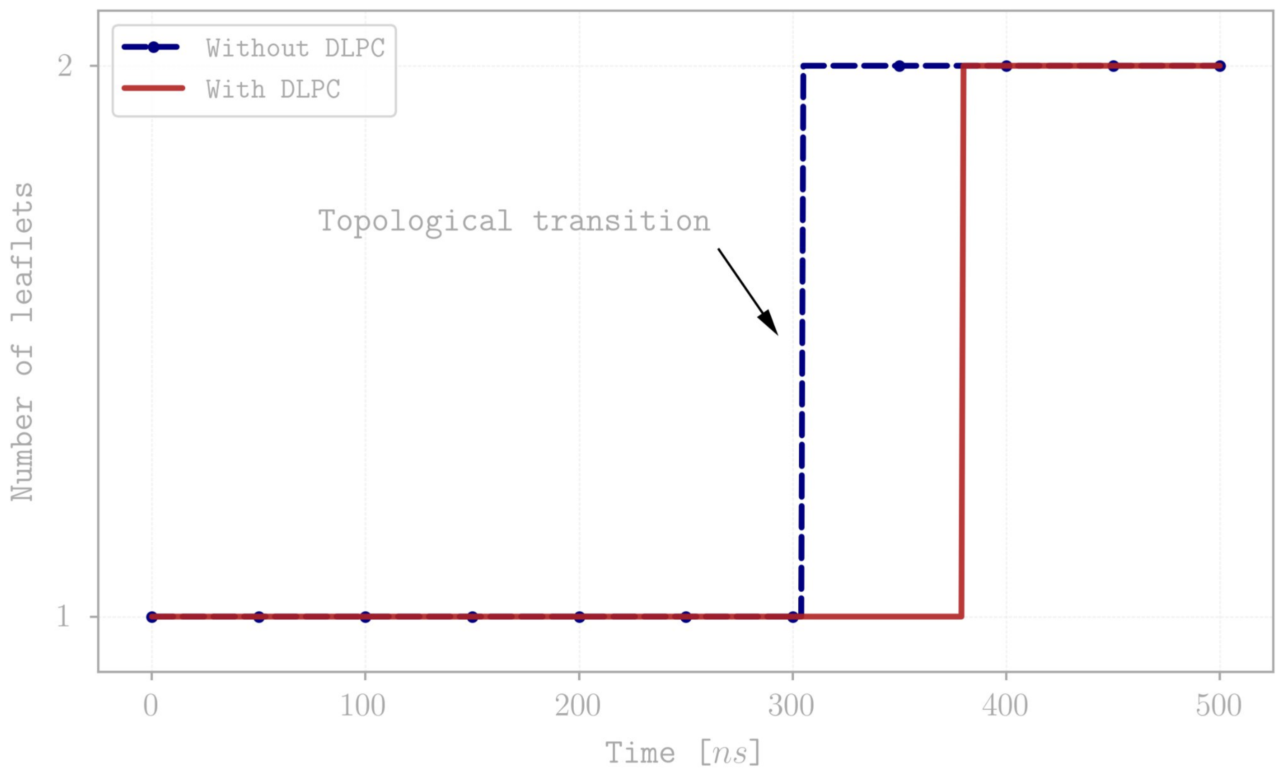


**Figure SF1:** The number of leaflets over time is shown for both Moebius membrane compositions. (WIP)

The diamond cubic phases were constructed using an approximation of the Schwarz diamond minimal surface[52]. Based on points sampled from this analytical approximation, triangulated meshes were created using a marching cubes algorithm[78]. Triangulated meshes were generated for both an individual unit cell and a 2×2×2 periodic array to demonstrate our ability to build larger system sizes of cubic phases[56].

The meshes were processed using the Pointillism subroutine to generate discretized point cloud representations. The constructed membrane consists of a pure monoolein lipid mixture, which was placed with an appropriate area per lipid of 0.375 $nm^2$. The systems were subsequently solvated and equilibrated following the standard protocol detailed above. Production simulations were extended to 2 microseconds to ensure sufficient sampling of phase dynamics.

To determine the stability of the cubic phase, we fitted the minimal surface mathematical model to the lipid positions from the simulation trajectories following the method described by Khelashvili et al.[52] Residual analysis from surface fits to 30 frames across the simulated trajectory showed a limited deviation from the theoretical minimal surface, confirming the preservation of the cubic structure throughout the simulation. Gaussian curvature on the fitted surface was calculated using the formula for implicitly defined surfaces, using the equation for the fitted surface previously described[56,79]. The results of this analysis are presented in Figure SX.

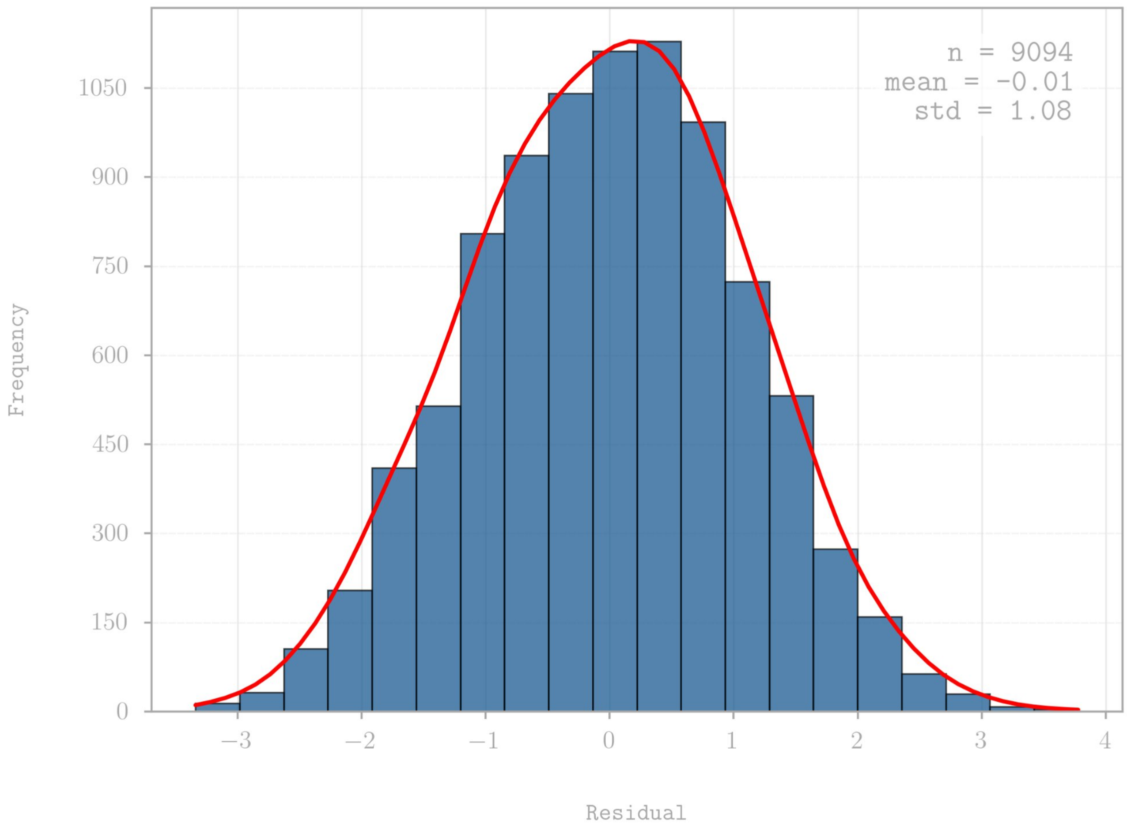


**Figure SF2:** Histogram of residuals from the fit of the diamond minimal surface to 30 frames from the trajectory of a 2x2x2 periodic system.

Furthermore, for the 2×2×2 periodic system, we performed a containment analysis to characterize the connectivity of the water channels. It is experimentally established that the diamond cubic phase contains two non-intersecting water channels. Using a voxel-based confinement analysis[57], we successfully identified these two distinct, non-intersecting water networks characteristic of the diamond cubic phase, further validating the stability of the constructed membrane system.

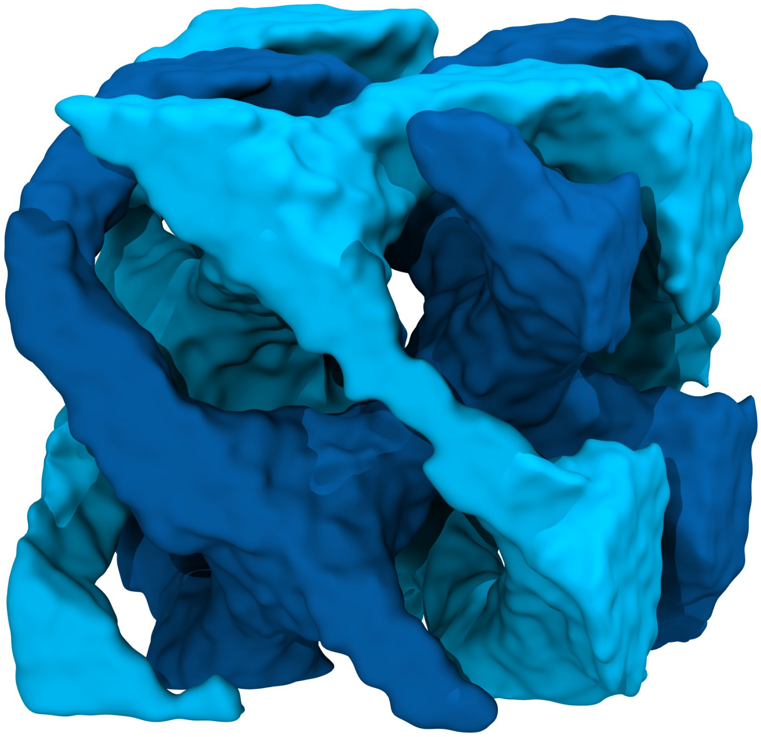

**Figure SF3:** Snapshot of the two non-intersecting water channels in the 2x2x2 lipid cubic phase (different channels are depicted by a different color).

##### Martini Globe

The created globe with the continents was subjected to simulation using Dry Martini. In multiple consecutive steps, the structure was minimized, and equilibrated.

The minimization consisted firstly of 1000 steps in double precision. The minimization was performed using the steepest descent algorithm.

As a second step, the globe was subjected to a series of two equilibration simulations, in which restraints were gradually turned off. All equilibration runs were conducted using the stochastic dynamics (sd) integrator at 310 K in an NVT ensemble. In the first run, the head groups were restrained with a timestep of 10 fs for 0.2 ns. In the second run, the restraints were lifted, the timestep was increased to 20 fs for 1.2 ns.

A final production simulation was conducted for 5 µs.

#### S5 The globe movie

See separate supplementary material. S5_globe_movie.mpg

The movie of the globe simulation has been reversed and a sky with stars has been placed in the background. The final five seconds show a plain image of the assembled continents. The video constitutes a non-scientific visualization of the case study and is mainly intended for outreach purposes.

#### S6 Non-Martini membranes

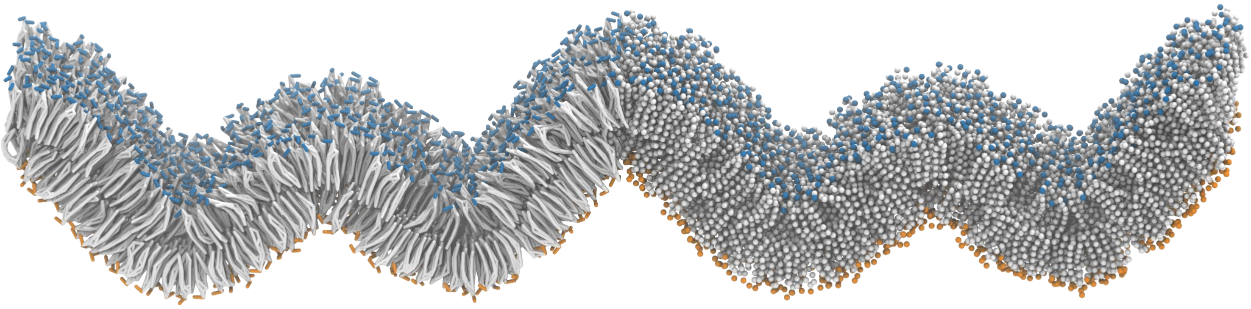

**Figure SF4: SIRAH and TS2CG** The figure shows an energy minimized Sirah POPC lipid membrane using the analytical shape function from TS2CG. In order to generate the structure, a custom LIB file was provided. The membrane is shown twice, once in a licorice representation showing the bonds and once showing the single beads. The head groups are colored blue or orange according to their monolayer.

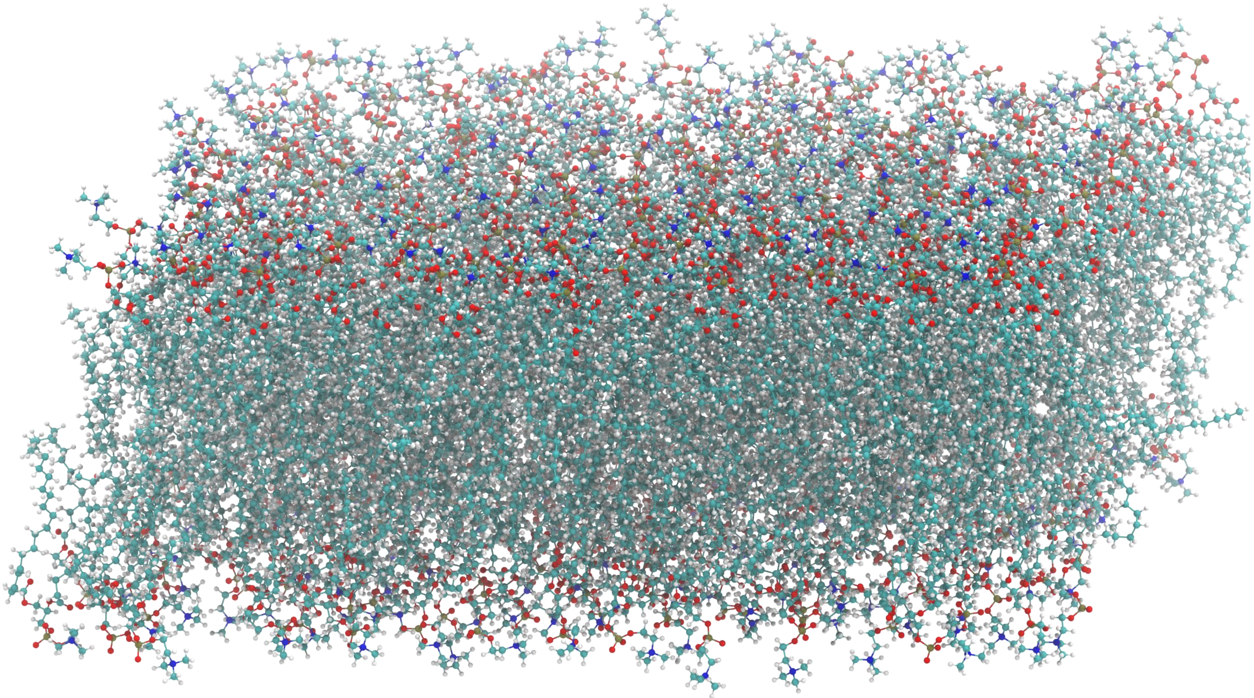

**Figure SF5: Charmm and TS2CG** The figure shows a POPC membrane in the CHARMM force field generated by TS2CG. A 10x10 nm flat membrane was built and the POPC lipids were subsequently minimized, solvated, and shortly equilibrated. Each atom is shown with nitrogens in blue, oxygens in red, carbons in cyan, phosphoruses in brown, and hydrogens in white.
