## Supplementary material for "TS2CG as a membrane builder": Details to S1

#### Tutorial 1: Creating a Simple Vesicle

The files needed for the tutorial can be manually downloaded here.

This tutorial will create a vesicle using a simple TS file (sphere). We chose this shape because it is small and can be easily run on your local machines. However, the same method applies to other TS files as well.

Start by selecting the **Sphere.tsi** file. Open it with a text editor to familiarize yourself with its structure. The **.tsi** file is an output file from DTS (Dynamically Triangulated Surface). It contains structural data about the simulation, including information on vertices, triangles, and the positions of proteins or other molecules. Specifically, the file includes:

1. **Vertices** represent points in space, typically defining the geometry of the simulated system. Each vertex includes its coordinates (x, y, z) and a domain identifier.
2. **Triangles** define the connections between vertices, forming the surfaces or mesh elements in the simulation. Each triangle references three vertices.
3. **Inclusions** represent proteins or other molecular entities, with information about their position, type, and associated vertex.

The first step in converting (backmapping) a TS file into a coarse-grained (CG) structure is to increase the number of vertices using a pointillism operation, which PLM performs. During this step, the two monolayers are also generated.

The command to run this step is:

```
TS2CG PLM -TSfile Sphere.tsi -bilayerThickness 3.8  -rescalefactor  4   4   4
```

This command reads the **.tsi** file and extends the structure based on the specified rescaling factors (Note: in the previous version, the rescalefactor had only one value). The output consists of two folders:

1. A folder containing visualization files (pointvisualization_data).
2. A folder that can be read by the CG Membrane Builder (PCG) script (further details provided in the following)

In the **pointvisualization_data** folder, you will find GROMACS-compatible structure files (**.gro**) for both the upper and lower monolayers, along with their respective topology files (**.top**). Additionally, you will find files compatible with common visualization software like ParaView and VMD. You can use **TS2CG PLM -h** for available flags.

The second step in creating a vesicle is to place lipids on the generated points using **PCG**. To do this, you'll need to create a **.str** file that defines the lipid composition for both monolayers. Using any text editor, create an **input.str** file and include the following content:

```
[Lipids List]

Domain 0
POPC  1 1 0.64
End
```

This specifies that your system will consist of a single lipid domain containing POPC in both the upper and lower monolayers with the same distribution ratio and an area per lipid (APL) of 0.64 $nm^2$ for POPC.

Next, we need a lipid structure file (**.LIB**). This file defines the lipid connectivity, which is used to position the lipid beads onto the points generated earlier. Although creating this file is straightforward, it can be time-consuming if you are working with many different lipids. Fortunately, we already have a **Martini3.LIB** file that contains all Martini3 lipids. You can find it in the files folder. Using these two files, you can now execute **PCG**:

```
TS2CG PCG -str input.str -Bondlength 0.2 -LLIB ./files/Martini3.LIB -defout system
```

The outputs will be **system.gro** (the structure file) and **system.top** (the topology file). These files contain the final vesicle configuration and are ready for further simulation or analysis.

```
;This file was generated by TS2CG membrane builder script i.e., PCG
 [ system ]
 Expect a large membrane
 [ molecules ]
; domain 0
 ;  in the upper monolayer
     POPC  6256
; domain 0
 ;  in the lower monolayer
     POPC  3876
```

This concludes the TS2CG part of the tutorial. Simulating the created vesicle would be the canonical next step. A script named **run_tut1.sh** is available in the **tut1** folder. This script will generate a POPC vesicle and run the TS2CG outputs using GROMACS:

To run the outputs from TS2CG, follow these steps:

1. **Energy Minimization with Softcore Potential**: Perform a short, 50-step energy minimization using the softcore potential, applying restraints to the lipid headgroups and protein backbones. Note that this step is optional and may not be necessary for all systems.
2. **Energy Minimization without Solvent**: Conduct a standard energy minimization, excluding solvent from the system.
3. **Short Equilibration without Solvent**: Run a brief equilibration step without solvent.

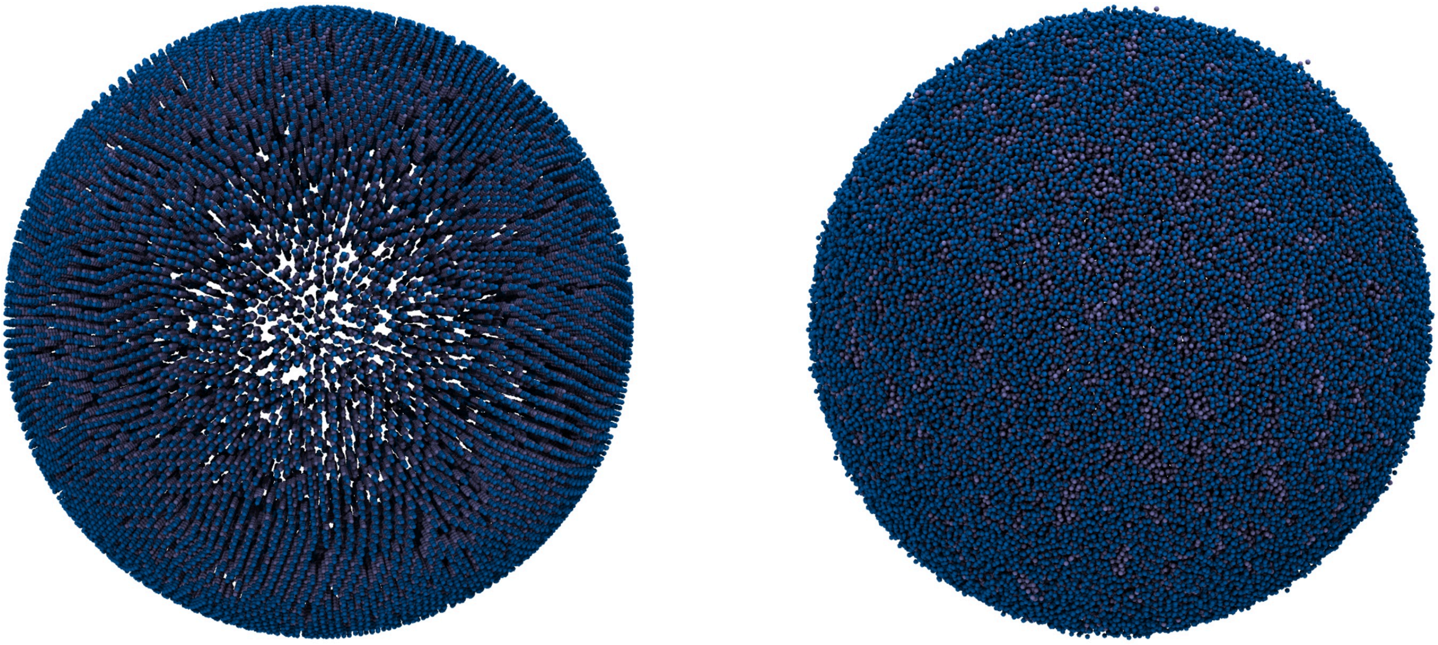

**Fig. 1.** Initial structure of the created POPC vesicle (left) and the vesicle after a brief vacuum simulation using GROMACS (right). Visualized with VMD.

#### Tutorial 2: Creating Lipid Mixture

The files needed for the tutorial can be manually downloaded here.

In this Tutorial, we are going to create a vesicle with a 50/50 mixture of POPC and DOPC in both monolayers, execute **PLM** using **Sphere.tsi** and **PCG** with the provided **.str** file as follows:

```
[Lipids List]

Domain 0
POPC  0.5 0.5 0.64
DOPC  0.5 0.5 0.67
End
```

```
TS2CG PLM -TSfile Sphere.tsi -bilayerThickness 3.8  -rescalefactor  4  4  4
```

```
TS2CG PCG -str input.str -Bondlength 0.2 -LLIB ./files/Martini3.LIB -defout system
```

The **PLM** tool reads the **.tsi** file and extends the structure according to the specified rescaling factors. This generates a set of points based on the desired system size and lipid distribution. The **PCG** tool places lipids on the generated points, producing two output files:

- **system.gro**: contains the coordinates and structure of the generated lipid system.
- **system.top**: the topology file, which should have the following format:

```
;This file was generated by TS2CG membrane builder script i.e., PCG
 [ system ]
 Expect a large membrane
 [ molecules ]
; domain 0
 ;  in the upper monolayer
     POPC  3056
     DOPC  3056
; domain 0
 ;  in the lower monolayer
     POPC  1893
     DOPC  1893
```

You can find a script named **run_tut2.sh** in the **tut2** folder. This script will generate a mixed POPC/DOPC vesicle and run the TS2CG outputs using GROMACS, following the steps below:

1. **Energy Minimization with Softcore Potential**: Perform a short, 50-step energy minimization using the softcore potential, applying restraints to the lipid headgroups and protein backbones. Note that this step is optional and may not be necessary for all systems.
2. **Energy Minimization without Solvent**: Conduct a standard energy minimization, excluding solvent from the system.
3. **Short Equilibration without Solvent**: Run a brief equilibration step without solvent.

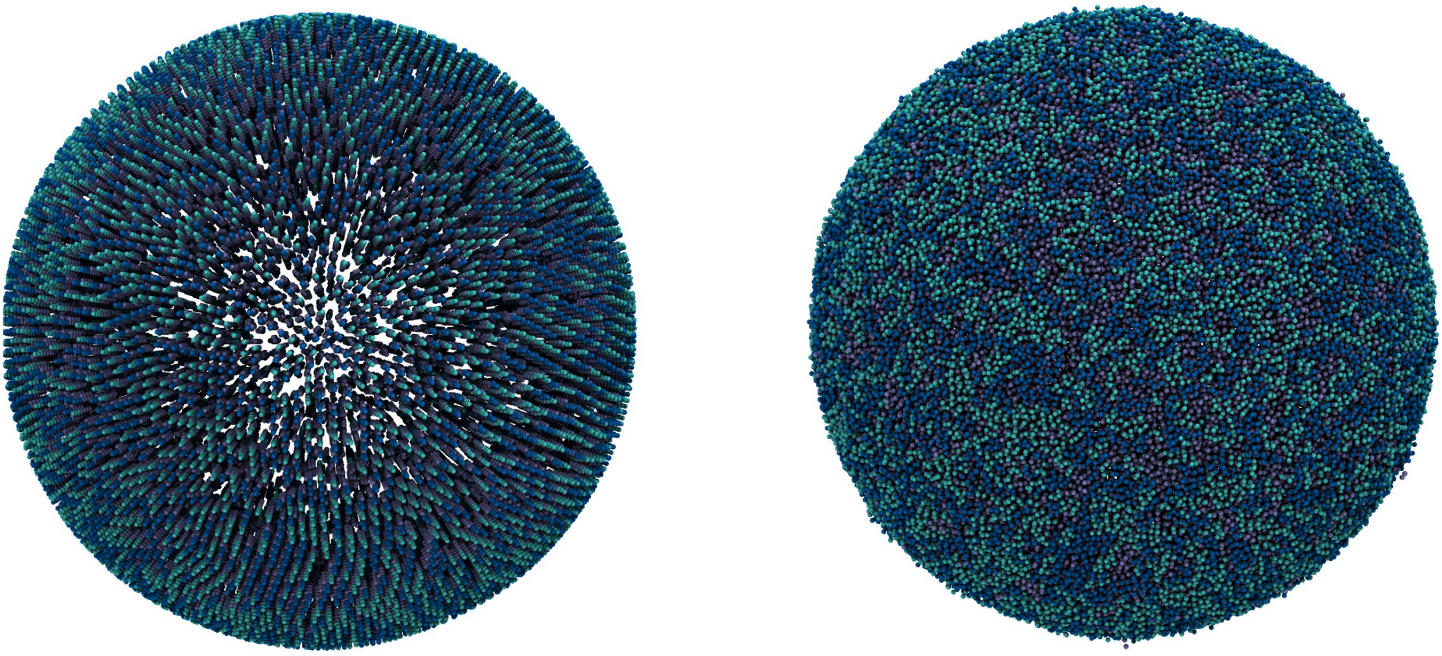

**Fig. 2.** Initial structure of the created mixed POPC/DOPC vesicle (left) and the vesicle after a brief vacuum simulation using GROMACS (right). Visualized with VMD.

#### Tutorial 3: Membrane Domains

The files needed for the tutorial can be manually downloaded here.

To create a membrane containing two or more distinct lipid domains, we need to modify the **.tsi** file. These changes can be made either manually or with the routines supplied in the **PointUpdaterClass (PUC)**. First, we will attempt the manual method, and then use the **DAI** program in **PUC** to automatize it.

#### 3-1. Manual Method:

First, we'll manually update the **.tsi** file. Open the **Sphere.tsi** file as a text file and make the following domain changes:

- Change lines 48, 112, 113, 117, and 124 to domain 1.
- Change lines 45, 74, 81, and 82 to domain 2.

Here's an example of a line from **Sphere.tsi** with the domain information added:

```
45     23.4753129908     28.0982228567     26.6006234584    2
```

The remaining lines remain unchanged as an empty domain field will default to domain 0.

---

> NOTE
>
> The line selections above are somewhat arbitrary. For a more precise choice of lines, you can use the following command:
>
> ```
> TS2CG PLM -TSfile Sphere.tsi -bilayerThickness 0 -rescalefactor 0.2 0.2 0.2 -Mashno 0
> ```
>
> This command will generate a down-scaled sphere while retaining the original number of vertices (using -Mashno 0) . Next, navigate to the **pointvisualization_data** folder and open **Upper.gro** in a visualization software of your choice (such as VMD). This will allow you to visualize the structure and identify specific lines to modify. Once you've selected the appropriate lines, update the **Sphere.tsi** file by adding the domain ID to the end of each selected line. Then, proceed with the tutorial.

---

Lastly, modify the **input.str** file to define the lipid composition for each domain:

```
[Lipids List]
Domain   0
POPC     1    1    0.64
End
Domain   1
DOPC     1    1    0.64
End
```

```
Domain   2
POPE     1    1    0.64
End
```

Next, execute **PLM** and **PCG** using the commands below to generate a vesicle with three lipid domains.

```
TS2CG PLM -TSfile Sphere.tsi -bilayerThickness 3.8  -rescalefactor  4    4    4
```

```
TS2CG PCG -str input.str -Bondlength 0.2 -LLIB ./files/Martini3.LIB -defout system
```

The outputs will be **system.gro** (the structure file) and **system.top** (the topology file). These files contain the final vesicle configuration and are ready for further simulation or analysis. The topology file includes the following content:

```
;This file was generated by TS2CG membrane builder script i.e., PCG
 [ system ]
 Expect a large membrane
 [ molecules ]
; domain 0
 ;  in the upper monolayer
     POPC  5834
; domain 0
 ;  in the lower monolayer
     POPC  3614
; domain 1
 ;  in the upper monolayer
     DOPC  259
; domain 1
 ;  in the lower monolayer
     DOPC  160
; domain 2
 ;  in the upper monolayer
     POPE  162
; domain 2
 ;  in the lower monolayer
     POPE  100
```

You can find a script named **run_tut3_1.sh** in the **tut3** folder. This script will generate a mixed POPC/DOPC/POPE vesicle with three domains and run the TS2CG outputs using GROMACS. It includes the following steps:

1. **Energy Minimization with Softcore Potential**: Perform a short, 50-step energy minimization using the softcore potential, applying restraints to the lipid headgroups and protein backbones. Note that this step is optional and may not be necessary for all systems.

2. **Energy Minimization without Solvent**: Conduct a standard energy minimization, excluding solvent from the system.
3. **Short Equilibration without Solvent**: Run a brief equilibration step without solvent.

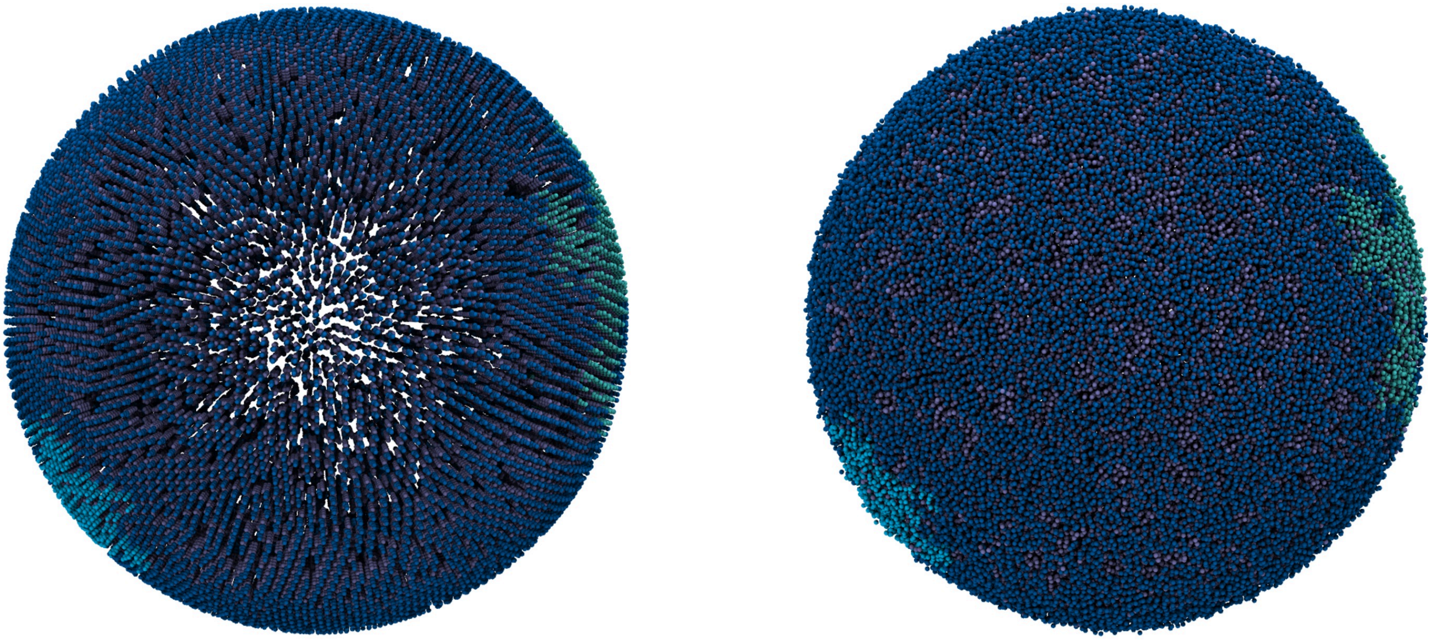

**Fig. 3-1.** Initial structure of the created vesicle with POPC/DOPC/POPE domains (left) and the vesicle after a brief vacuum simulation using GROMACS (right). Visualized with VMD.

#### 3-2. Automatized Circular Domains:

An alternate approach to place circular domains at certain positions is provided with the **DAI** tool. Consider **Sphere_2.tsi**, which is an exact copy of the original **Sphere.tsi**.

Use **Sphere_2.tsi** without adding any lipid domains and execute **PLM** with the following command:

```
TS2CG PLM -TSfile Sphere_2.tsi -bilayerThickness 3.8 -rescalefactor 4   4   4
```

Next, identify the beads on which you want to create lipid domains. In this example, points [5, 22] are selected for domain 1 and point [30] for domain 2, while all other points remain in domain 0.

To create a circular lipid domain with domain ID 1 and an example radius of 4 around points [5, 22], execute the following command:

```
TS2CG DAI -p point -r 4 -d 1 -dummy 5,22
```

r specifies the radius of the circular domain around the points, and d tells the DAI program to populate it with domain 1.

Similarly, to create a circular domain for domain 2:

```
TS2CG DAI -p point -r 5 -d 2 -dummy 30
```

This will update the point folder and place a circular domain with domain ID 2 and a radius of 5 around point 30. Note that if the circle for domain 2 overlaps a circle for domain 1, it will overwrite domain 1.

After defining the domains, place the lipids using the **.str** file with the following command:

```
TS2CG PCG -str input.str -Bondlength 0.2 -LLIB ./files/Martini3.LIB -defout system
```

This will generate the final coarse-grained structure, incorporating the lipid domains defined in the previous steps. It can be simulated as shown in the first part of this tutorial (**run_tut3_2.sh**).

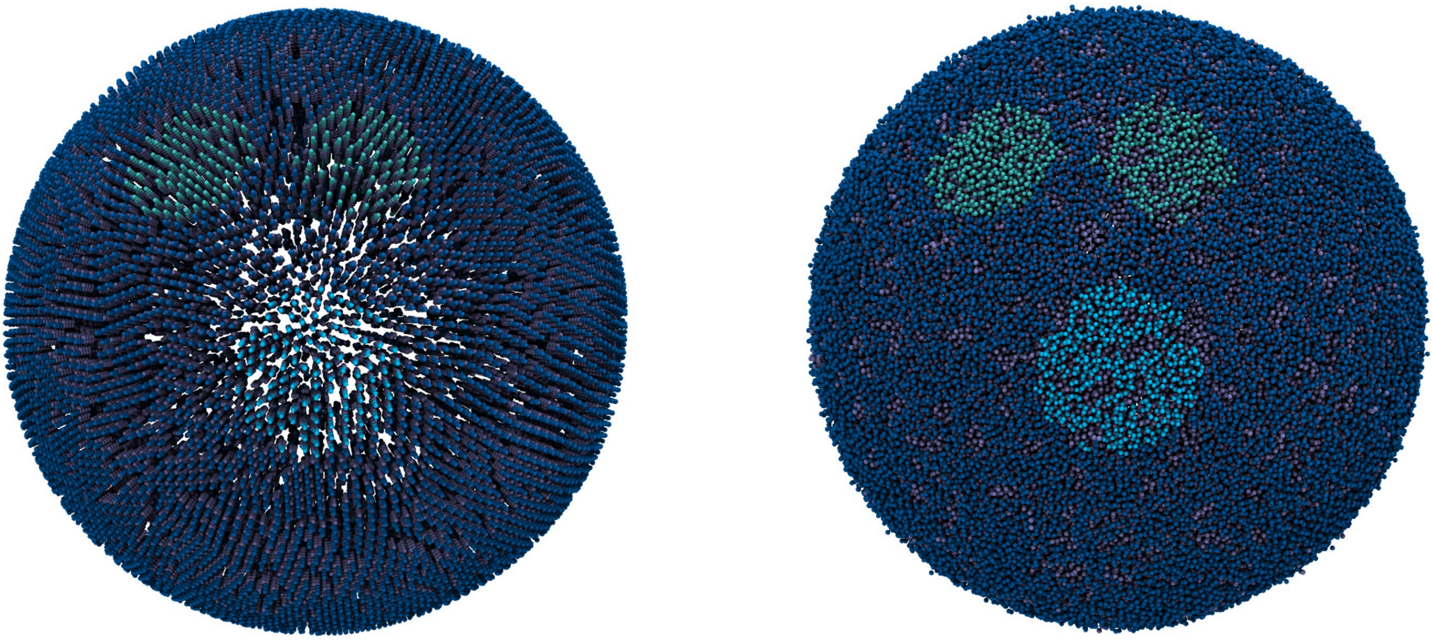

**Fig. 3-2.** Initial structure of the created vesicle with POPC/DOPC/POPE circular domains (left) and the vesicle after a brief vacuum simulation using GROMACS (right). Visualized with VMD.

#### Tutorial 4: Adding Protein to the Membrane

The files needed for the tutorial can be manually downloaded here.

In this tutorial, we will add two types of proteins to a vesicle containing POPC lipids. The proteins are named **P1.gro** and **P2.gro**, which will be referred to as **protein1** and **protein2** for the remainder of this tutorial.

First, we need to select vertices for protein placement in the **.tsi** file. Afterward, we will match their corresponding names in the **.gro** file with those in the **.str** file. To achieve this, we will perform some modifications on the **.tsi** and **.str** files.

Start by using the command below to obtain a **.gro** file containing the vertex positions of our vesicle:

```
TS2CG PLM -TSfile Sphere.tsi -bilayerThickness 0  -rescalefactor 0.2 0.2 0.2 -Mashno 0
```

Identify the vertices where you would like to place the proteins by opening the **Upper.gro** file from the **pointvisualization_data** folder in a visualization software of your choice (such as VMD). For this tutorial, we selected vertices **5** and **22** to place two copies of **protein1**, and vertex **30** for one copy of **protein2**.

Open the **Sphere.tsi** file using a text editor and scroll to the bottom to find the inclusion section. Since we want to add three proteins, change the existing **0** to **3**. On the next line, add the protein information. For each protein, you will need to provide three integer values and two float values:

1. **Protein Index**: This should start from **0**.
2. **Protein Type ID**: Use **1** for **protein1** and **2** for **protein2** (the ID can be any number, but it must match the one in **input.str**, as explained below).
3. **Vertex Index**: This is the index of the vertex where the protein will be placed.
4. **Orientation**: The last two numbers represent the orientation of the protein in the local coordinate frame of the vertex; these should be a unit two-dimensional vector.

The inclusion section of the **Sphere.tsi** file should look like this:

```
inclusion        3
0   1   5    0   1
1   1   22   0   1
2   2   30   0   1
```

---

> NOTE
>
> In this tutorial, **protein2** (**P2.gro**) represents VDAC1, or voltage-dependent anion channel 1, which is

a pore-forming protein. For proteins like this, we need to ensure that lipid molecules do not occupy the space within the channel pore. To achieve this, we will add an exclusion section to the **.tsi** file.

> Now, open the **Sphere.tsi** file in a text editor, and add the exclusion section at the end of the file. Since there is only one **protein2** in the system that should be free of lipids, write **1** in front of the exclusion line.
>
> For each pore that you want to create, you will need to provide three values:
>
> 1. **Pore Index**: Start from **0**.
> 2. **Vertex Index**: The index of the vertex where we want to remove the points
> 3. **Pore Radius**: This is the radius of the pore that we want to create around the specified vertex. The exclusion section of the **Sphere.tsi** file should look like this:

> ```
> exclusion    1
> 0     30     1
> ```

---

Now, open **P1.gro** and **P2.gro** from the **files** folder and change the first line to reflect the names of the proteins: **protein1** and **protein2**. These files should be included at the top of your input.str file as follows:

```
include P1.gro
include P2.gro
```

The final step is to define the proteins in the **input.str** file. In addition to including the protein **.gro** file names in the header, you also need to provide information about the protein placement:

```
[Protein List]
protein1     1     0.01     0     0     -2.5
protein2     2     0.01     0     0     -2.5
End Protein
```

The first and last lines serve as markers to indicate the beginning and end of protein definitions. The lines in between correspond to the number of unique proteins being defined.

In this example, there are two unique proteins, so there are two intermediate lines.

Each intermediate line contains three key pieces of information:

1. The first entry is the protein name as listed in the **.gro** file.
2. The second entry is the protein type ID, which corresponds to the ID used in the inclusion section of the **.tsi** file.
3. The last entry specifies how much the proteins should be shifted in the normal direction relative to the membrane surface.

The remaining three numbers are not utilized in the current approach.

So, the **input.str** file will look as follows:

```
include  P1.gro
include  P2.gro
[Lipids List]
Domain  0
; lipidname ratio_up ratio_down  APL
POPC    1    1    0.64
End
[Protein List]
```

```
protein1     1     0.01     0     0     -2.5
protein2     2     0.01     0     0     -2.5
End Protein
```

Finally, execute **PLM** and **PCG** using the commands below (or as shown in the previous tutorials). The result should be a POPC membrane with three proteins.

```
TS2CG PLM -TSfile Sphere.tsi -bilayerThickness 3.8  -rescalefactor 4 4 4
```

```
TS2CG PCG -str input.str -Bondlength 0.2 -LLIB ./files/Martini3.LIB -defout system
```

A script named **run_tut4.sh** is available in the **tut4** folder. This script will generate a POPC vesicle containing three proteins (two copies of **protein1** and one copy of **protein2**) and run the TS2CG outputs using GROMACS:

1. **Energy Minimization without Solvent**: Conduct a standard energy minimization, excluding solvent from the system.
2. **Short Equilibration without Solvent**: Run a brief equilibration step without solvent.

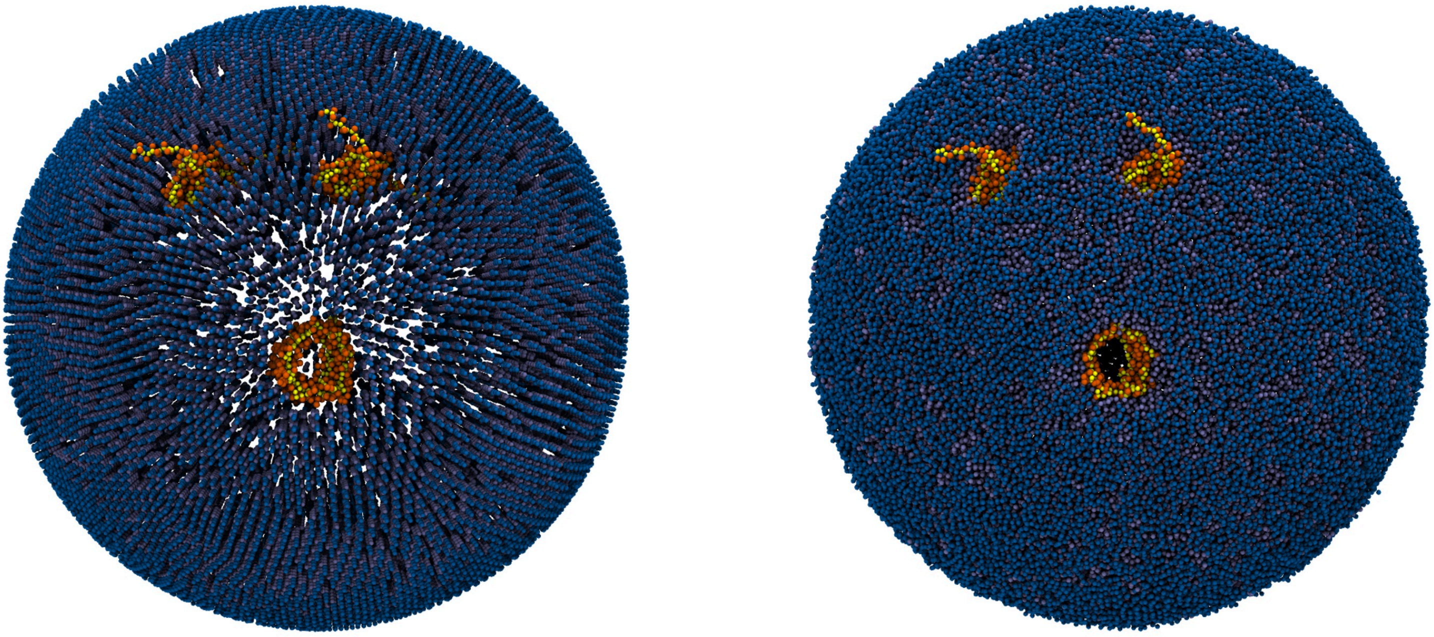

**Fig. 4.** Initial structure of the created POPC vesicle containing three proteins (left) and the vesicle after a brief vacuum simulation using GROMACS (right). Visualized with VMD.

#### Tutorial 5: Protein with a Specific Domain

The files needed for the tutorial can be manually downloaded here.

In this tutorial, we will add two types of proteins to a vesicle and define a specific lipid domain for each protein copy. The proteins are named **P1.gro** and **P2.gro**, which will be referred to as **protein1** and **protein2** for the remainder of this tutorial.

To do this, we need to modify the **.tsi** file either manually or with the routines supplied in the **PointUpdaterClass (PUC)**. First, we will attempt the manual method, and then use the **DAI** program in **PUC** to automatize it.

#### 5-1 Manual Method:

First, we need to select vertices for protein placement in the **.tsi** file. Afterward, we will match their corresponding names in the **.gro** file with those in the **.str** file. To achieve this, we will perform some modifications on the **.tsi** and **.str** files.

Start by using the command below to obtain a **.gro** file containing the vertex positions of our vesicle:

```
TS2CG PLM -TSfile Sphere.tsi -bilayerThickness 0  -rescalefactor 0.2 0.2 0.2 -Mashno 0
```

Identify the vertices where you would like to place the proteins by opening the **Upper.gro** file (located in the **pointvisualization_data** folder) in a visualization software of your choice, such as VMD. For this tutorial, we selected vertices **5** and **22** to place two copies of **protein1**, and vertex **30** for one copy of **protein2**.

Open the **Sphere.tsi** file using a text editor and scroll to the bottom to find the inclusion section. Since we want to add three proteins, change the existing **0** to **3**. On the next line, add the protein information. For each protein, you will need to provide three integer values and two float values:

1. **Protein Index**: This should start from **0**.
2. **Protein Type ID**: Use **1** for **protein1** and **2** for **protein2** (the ID can be any number, but it must match the one in **input.str**, as explained below).
3. **Vertex Index**: This is the index of the vertex where the protein will be placed.
4. **Orientation**: The last two numbers represent the orientation of the protein in the local coordinate frame of the vertex; these should be a unit two-dimensional vector.

The inclusion section of the **Sphere.tsi** file should looks as follows:

```
inclusion       3
0    1    5     0    1
1    1    22    0    1
2    2    30    0    1
```

Now, let's assign specific lipid domain IDs to the vertices where the proteins are located in the **Sphere.tsi** file. Since each protein at vertices 5, 22, and 30 will be surrounded by a specific lipid type, we need to update these lines with corresponding lipid domain IDs:

- **Vertex 5** and **vertex 22**: Change these lines to **domain 1** to assign the first lipid type.
- **Vertex 30**: Change this line to **domain 2** to assign the second lipid type.

For example, the line for vertex 5 should look like this after adding the lipid domain ID:

```
5     22.0396876425     23.6080597437     26.8858740866    1
```

---

> NOTE
>
> In this tutorial, **protein2** (**P2.gro**) represents VDAC1, or voltage-dependent anion channel 1, which is a pore-forming protein. For proteins like this, we need to ensure that lipid molecules do not occupy the space within the channel pore. To achieve this, we will add an exclusion section to the **.tsi** file.
>
> Now, open the **Sphere.tsi** file in a text editor, and add the exclusion section at the end of the file. Since there is only one **protein2** in the system that should be free of lipids, write **1** in front of the exclusion line.
>
> For each pore that you want to create, you will need to provide three values:
>
> 1. **Pore Index**: Start from **0**.
> 2. **Vertex Index**: The index of the vertex where we want to remove the points
> 3. **Pore Radius**: This is the radius of the pore that we want to create around the specified vertex. The exclusion section of the **Sphere.tsi** file should look like this:
>
> ```
> exclusion    1
> 0     30     1
> ```

---

To complete the setup, follow these steps:

1. Update Protein Names in **.gro** Files:
    - Open **P1.gro** and **P2.gro**.
    - Change the first line in each file to match the desired protein names: **protein1** for **P1.gro** and **protein2** for **P2.gro**.
2. Include Protein Files in the **input.str**:
    - Add the names of the **.gro** files for each protein at the top of the **input.str** file.
3. Define the proteins in the **input.str**:
    - The [Protein List] section begins and ends with header and footer lines to mark the section.
    - Each unique protein should have a line with three main pieces of information:

        - **Protein Name**: This should match the name in the **.gro** file (e.g., protein1, protein2).
        - **Protein Type ID**: Matches the ID specified for the protein in the inclusion section of the **.tsi** file.
        - **Position Shift**: Specifies the offset of the protein in the normal direction relative to the membrane surface.
    - The remaining three numbers on each line are placeholders and are not used in the current approach.
4. Modify the Lipid Section:
    - In the [Lipids List] section, specify each lipid domain, the lipid type, and the associated properties as shown in the example below.

Here's the final format of the **input.str** file:

```
include P1.gro
include P2.gro

[Lipids List]
Domain   0
POPC     1   1   0.64
End
Domain   1
DOPC     1   1   0.64
End
Domain   2
POPE     1   1   0.64
End

[Protein List]
protein1     1   0.01   0   0   -2.5
protein2     2   0.01   0   0   -2.5
End Protein
```

This configuration ensures that each protein and lipid domain is correctly defined for simulation, with proteins positioned at specified vertices and surrounded by their designated lipid domains.

Now perform **PLM** and **PCG**:

```
TS2CG PLM -TSfile Sphere.tsi -bilayerThickness 3.8  -rescalefactor 4 4 4
```

```
TS2CG PCG -str input.str -Bondlength 0.2 -LLIB ./files/Martini3.LIB -defout system
```

You can find a script named **run_tut5_1.sh** in the **tut5** folder. This script will generate a vesicle with three lipid domains (POPC, DOPC, and POPE) and three proteins. Each of the three proteins is positioned within a specific lipid domain. It will also run the TS2CG outputs in GROMACS, following these steps:

1. **Energy Minimization without Solvent**: Conduct a standard energy minimization, excluding solvent from the system.
2. **Short Equilibration without Solvent**: Run a brief equilibration step without solvent.

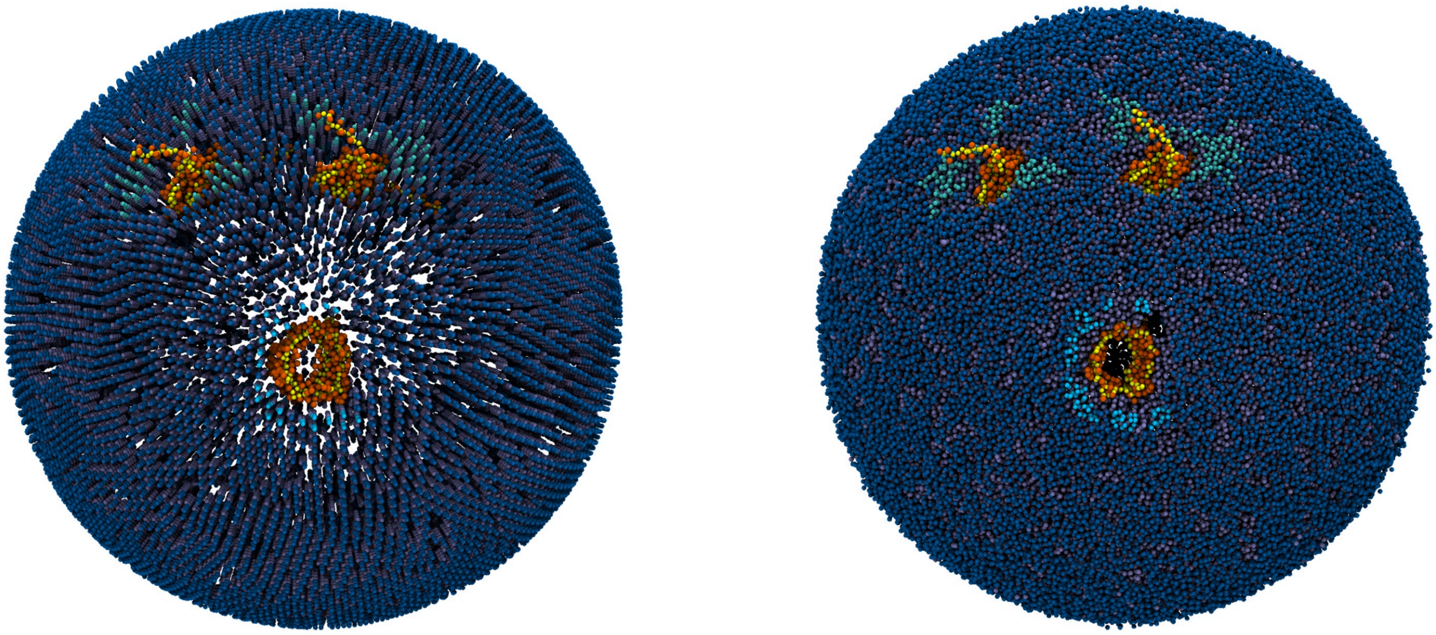

**Fig. 5-1.** Initial structure of the created vesicle with POPC/DOPC/POPE domains and three proteins (left) and the vesicle after a brief vacuum simulation using GROMACS (right). Visualized with VMD.

##### 5-2. Automatized Circular Domains:

As demonstrated in the automatized section of tutorial 3, you can add circular domains around each included protein using the **DAI** command. To do this, refer to **Sphere_2.tsi**, which contains the protein inclusions without lipid domains. Setting lipid domains manually is not necessary, as the circular domains can be configured automatically around each protein.

The inclusion and exclusion sections in **Sphere_2.tsi** looks like:

```
inclusion       3
0    1    5     0    1
1    1    22    0    1
2    2    30    0    1

exclusion  1
0    30    1
```

First, execute **PLM**:

```
TS2CG PLM -TSfile Sphere_2.tsi -bilayerThickness 3.8  -rescalefactor 4 4 4
```

To place a circular lipid domain with domain ID 1 and a radius of 4 around protein1, execute the following commands (r specifies the radius, d the domain, and T the inclusion type as specified in the inclusions):

```
TS2CG DAI -p point -r 4 -d 1 -t 1
```

For domain 2 around protein2, execute the command again:

```
TS2CG DAI -p point -r 4 -d 2 -t 2
```

This will update the point folder and place a circular domain with domain ID 2 and a radius of 4 around protein2.

**Note**, it might happen, that the circular domains overlap; in that case, the second command will overwrite the domain specification of the first.

Next, execute **PCG** to position the lipids and proteins in their specified locations:

```
TS2CG PCG -str input.str -Bondlength 0.2 -LLIB ./files/Martini3.LIB -defout system
```

The outputs generated by TS2CG can be simulated using GROMACS, following the steps outlined in the first part of this tutorial (**run_tut5_2.sh**).

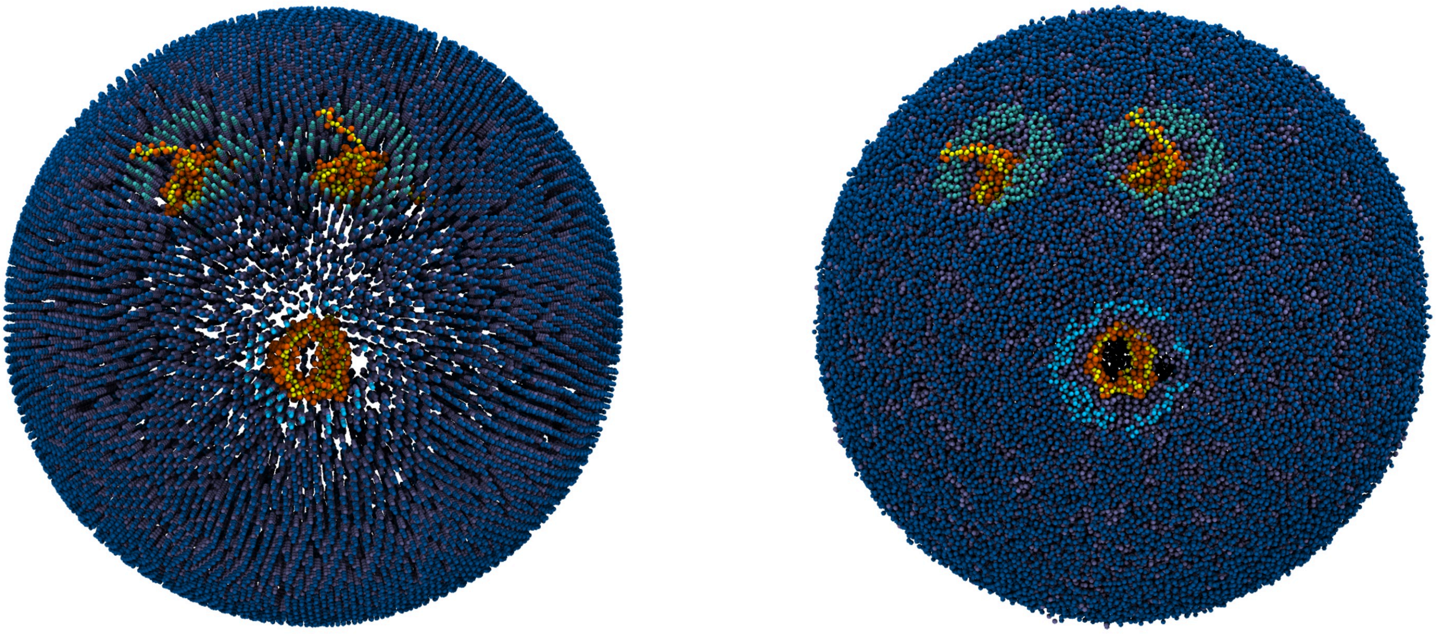

**Fig. 5-2.** Initial structure of the created vesicle with POPC/DOPC/POPE circular domains and three proteins (left) and the vesicle after a brief vacuum simulation using GROMACS (right). Visualized with VMD.

### Tutorial 6: Fixed Shapes

The files needed for the tutorial can be manually downloaded here.

**PCG** enables the generation of membranes with specific shapes, including flat bilayers, spheres, cylinders, and other periodic structures, which can be defined as a sum of 1D Fourier modes. This process does not require a **.tsi** file. In the **.str** file, shape information is defined as follows to create different structures.

```
Cylinder                                    1D Fourier Shape
[Shape Data]                                [Shape Data]
ShapeType    Cylinder                       ShapeType    1D_PBC_Fourier
Box    40    40    40                       Box    30    10    20
Density      2                              Density      3     1
Thickness    3.8                            Thickness    3.8
Radius       12                             WallRange    0     1    0    1
End                                         Mode         1.5   1    0
                                            Mode         2.5   2    0

Sphere                                      Flat
[Shape Data]                                [Shape Data]
ShapeType    Sphere                         ShapeType    Flat
Box    40    40    40                       Box    40    40    40
Density      2                              Density      2     2
Thickness    3.8                            Thickness    3.8
WallDensity  1       1                      WallRange    0     1    0    1
DL           0.2                            End
Radius       15
End
```

**Note**: While the shape type must be specified, the other options are optional. You can use default values for the remaining parameters.

##### 6-1. 1D Fourier-shaped POPC bilayer

To create a 1D Fourier shape POPC bilayer, you can use the following **.str** file as an example:

```
[Lipids List]
Domain      0
POPC        1   1   0.64
End

[Shape Data]
ShapeType   1D_PBC_Fourier
Box         30     10     20
WallRange   0      1      0      1
Density     3      1
Thickness   3.8
```

```
Mode        1.5    1     0
Mode        2.5    2     0
End
```

Using this file, you can now execute **PCG**:

```
TS2CG PCG -str input.str -Bondlength 0.2 -LLIB ./files/Martini3.LIB -defout system -function a
```

The outputs will be **system.gro** (the structure file) and **system.top** (the topology file). These files contain the final configuration of a 1D Fourier-shaped POPC bilayer, prepared for further simulation or analysis. Here's an example of what **system.top** looks like:

```
;This file was generated by TS2CG membrane builder script i.e., PCG
 [ system ]
 Expect a large membrane
 [ molecules ]
; domain 0
 ;  in the upper monolayer
     POPC  585
; domain 0
 ;  in the lower monolayer
     POPC  585
```

In the **tut6** folder, there is a script named **run_tut6_1.sh**. This script generates a 1D Fourier-shaped POPC membrane and runs the TS2CG outputs using GROMACS with the following steps:

1. **Energy Minimization with Softcore Potential**: Perform a short, 50-step energy minimization using the softcore potential, applying restraints to the lipid headgroups and protein backbones. Note that this step is optional and may not be necessary for all systems.

2. **Energy Minimization without Solvent**: Conduct a standard energy minimization, excluding solvent from the system.

3. **Short Equilibration without Solvent**: Run a brief equilibration step without solvent.

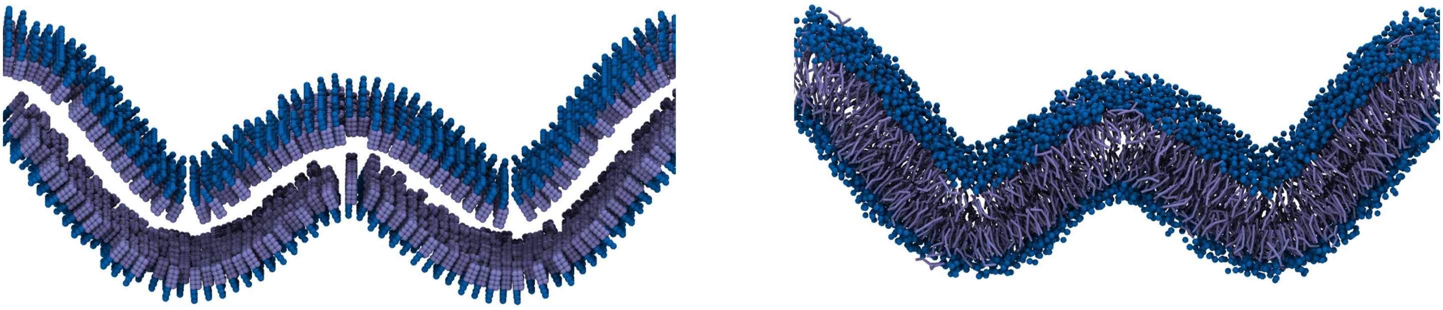

**Fig. 6-1.** Initial structure of the 1D Fourier-shaped POPC membrane (left) and the system after a brief vacuum simulation using GROMACS (right). Visualized with VMD.

##### 6-2. Creating the Wall

To generate wall beads, include the following flags in the PCG command line:

```
-Wall -WallBName WL
```

In addition to **.top** and **.gro** files, executing PCG with the above flag will also generate **Wall.itp** and **Wall.pdb** files. Note that it is up to you to to define the interaction between the wall beads and the bilayer to preserve the bilayer's shape. A recommended approach is to implement a repulsive Lennard-Jones (LJ) interaction between the wall beads and the lipid tail beads while ensuring that the wall beads remain invisible to other beads in the system. This system can be simulated using the steps described in the first part of this tutorial. During simulations, the wall beads should remain fixed in place. This can be achieved by either: 1. Using position_restraints as specified in the **Wall.itp** file, or 2. Defining a freeze group in the **.mdp** file. An example **.mdp** file is available in **files/mdp/Wall**. In this tutorial, the second method is used.

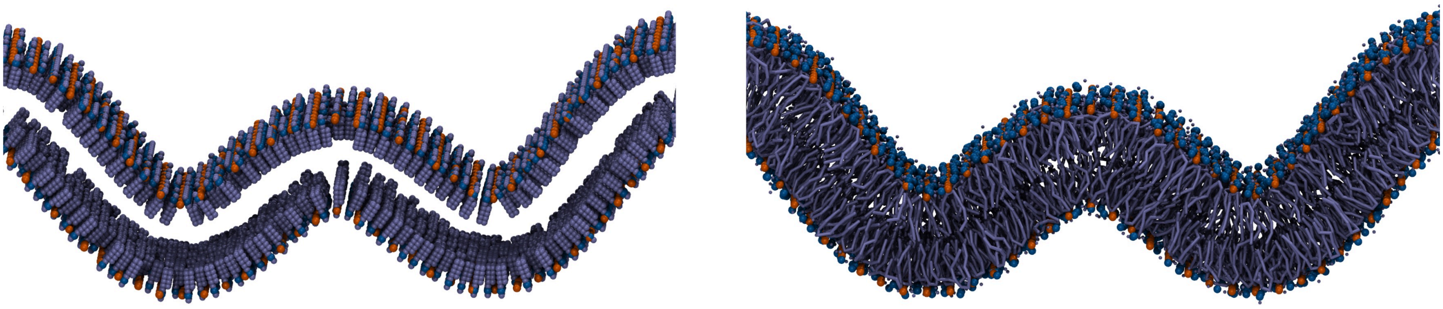

**Fig. 6-2.** Initial structure of the 1D Fourier-shaped POPC membrane with wall (left) and the system after a brief vacuum simulation using GROMACS (right). Visualized with VMD.

### Tutorial 7: Placing Lipids Based on Favored Curvature

The files needed for the tutorial can be manually downloaded here.

In this tutorial, we will create a curved lipid bilayer containing POPC and cardiolipin (CDL0), where the lipids will be placed based on their preferred curvature, bringing the output of TS2CG closer to equilibrium. The following equation is implemented into **TS2CG DOP**, allowing us to determine lipid placement by setting values for $C_0$ and $k$:

$e^{-k(C_1+C_2-C_0)^2}$

Here, $C_0$ represents the initial curvature, which is defined for each lipid type, while $k$ is the same for all lipid types. For instance, it is known that cardiolipin favors negatively curved regions. By selecting a $C_0$ value for cardiolipin that makes the equation negative, TS2CG will position cardiolipin in the negatively curved areas of the membrane. A higher $k$ value further enhances the precision of lipid placement. The lipid placement process involves three key steps:

1. Creating a point folder using **PCG**.

2. Modifying the point folder using **DOP**.
3. Building the lipid bilayer with **PCG**.

**Step 1. Create a point folder using PCG.**

Creating a point folder with **PCG** requires a **.str** file in the working directory. The **-WPointDir** flag in **PCG** generates the point folder. In this tutorial, we utilize a 1D Fourier shape. The provided **.str** file for this tutorial looks as follows:

```
[Lipids List]
;lipidnname ratio_up ratio_down  APL
POPC   1    1    0.64   ;Not necessary in this step but shouldn´t be removed.
End

[Shape Data]
ShapeType 1D_PBC_Fourier
Box    30    10    20
WallRange    0     1    0    1
Density      3     1
Thickness    4
Mode        1.5    1    0
Mode        2.5    2    0
End
```

Start by using the command below to obtain a **point** folder containing two files called **InnerBM.dat** and **OuterBM.dat** for our 1D_Fourier shape:

```
TS2CG PCG -str input.str -function analytical_shape -defout system -WPointDir
```

**Step 2. Modify the point folder using DOP**

In this step, we aim to modify the point folder from the previous step so that PCG (in the next step) knows where to place each lipid. Modifying the point folder requires a domain_input.txt file, which is provided. However, you can also use the following command to check the expected format and try creating your own:

```
TS2CG DOP -h
```

The domain_input.txt specifies which lipids to use, their respective proportions, C0 values for each lipid, and the area per lipid. In this tutorial, a 7:3 ratio of POPC to CDL0 is used, with $C_0$ = 0 for POPC and $C_0$ = -0.5 for CDL0.

```
; domain lipid percentage c0 density
0    POPC    .7    0.0    0.64
```

```
1    CDL0    .3   -0.5    1.2
```

Now, you only need to define one more variable: $k$, which is specified using a flag in the **TS2CG DOP** command. In this tutorial, we suggest setting the initial $k$ value to 10, but we strongly recommend experimenting with other values of $k$ to better understand its impact on lipid placement.

To modify the point folder, use the following command:

```
TS2CG DOP -i domain_input.txt -ni input_DOP.str -k 10
```

This command will generate a new, updated point folder and a new **.str** file called **input_DOP.str**, which contains information about the different domains based on the **domain_input.txt** file.

**Step 3. Build a bilayer based on the modified point folder using PCG**

In this final step, you will use **PCG** as done in previous tutorials, but with the files provided by **DOP** in the last step, the modified point folder, and the **input_DOP.str** file.

Execute the following **PCG** command to build the membrane:

```
TS2CG PCG -dts point -str input_DOP.str -LLIB ./files/Martini3.LIB -defout system
```

This will generate **system.top** and **system.gro** files. The **system.top** file can be visualized using VMD or any other preferred visualization software. The cardiolipin (CDL0) lipids should now be positioned in the negatively curved regions of the membrane, as shown in the image below, where CDL0 is represented in magenta.

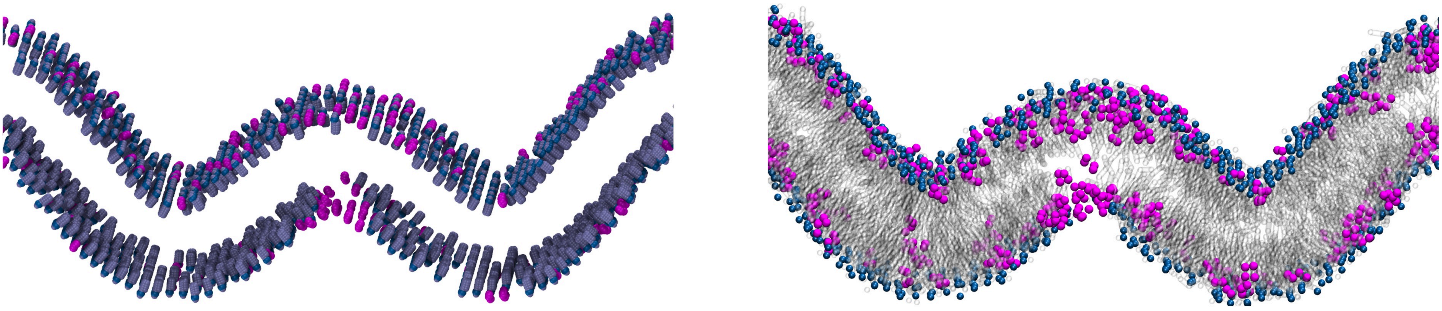

**Fig. 7.** Cardiolipin (CDL0) lipids (magenta) localized in negatively curved membrane regions

> NOTE:
>
> Finding a good $C_0$ value is a matter of trial and error or more rigorous studies into the distribution of lipids in a membrane. When you move on to your own systems, test various $C_0$ and in combination with different values of $k$ until you obtain the desired outcome.

> Furthermore, if you wish to compare different outcomes, remember to change the name of the new **.str**, **system.gro** and **system.top** since **TS2CG DOP** will overwrite them otherwise.
>
> ---

> ---
>
> NOTE:
>
> A negative $C_0$ will make Equation 1 negative for CDL0, instructing **PCG** to place CDL0 in the negatively curved regions, which is enhanced by a $k$-value of 10. Conversely, a positive $C0$ will have the opposite effect for POPC in this case.
>
> ---

A script named **run_tut7.sh** is available in the **tut7** folder. This script will generate a mixed membrane of POPC and CDL0. The membrane will be curved, and CDL0 will be placed in the negatively curved regions. The script will run the TS2CG output using GROMACS with the following steps.

1. **Energy Minimization with Softcore Potential**: Perform a short, 50-step energy minimization using the softcore potential, applying restraints to the lipid headgroups and protein backbones. Note that this step is optional and may not be necessary for all systems.

2. **Energy Minimization without Solvent**: Conduct a standard energy minimization, excluding solvent from the system.

3. **Short Equilibration without Solvent**: Run a brief equilibration step without solvent.

#### Tutorial 8: Add More Proteins and Place Proteins based on Curvature

The files needed for the tutorial can be manually downloaded here.

This tutorial is divided into two parts. In the first part you will use the input files from tutorial 4 to build a system and easily add more proteins to the system. Whereas, in the second part you build a curved membrane and place the proteins based on curvature (similarly to tutorial 7).

###### 8-1 Add more proteins

**Step 1: Generate point folder** Use the following command to generate the point folder.

```
TS2CG PLM -TSfile Sphere.tsi -bilayerThickness 3.8 -rescalefactor 4 4 4  -o point_vesicle
```

Since you are using the input files from Tutorial 4, the output from **PLM** will contain three proteins in total, two of type 1 and one protein of type 2. This can be seen in **point_vesicle/IncData.dat**.

```
< Inclusion NoInc       3   >
< id typeid pointid lx ly lz  >
```

```
          0      1      5       0.499      -0.864      0.059
          1      1      22      0.481      -0.765     -0.429
          2      2      30     -0.177      -0.945      0.275
```

**Step 2: Add more proteins**

In tutorial 4 you would now use **TS2CG PCG** to build the system but in this tutorial, you include one more step, **TS2CG INU**. Use **TS2CG INU -h** for available flags. This function will modify the point folder by adding more proteins of your choice. For example, to add three proteins of type 1 in the outer leaflet you can run the following command:

```
TS2CG INU -p point_vesicle -n 3 -t 1 -r 5 -o point_vesicle_new -l outer
```

This command will modify the **point** folder and save the updated **point** folder as **point_vesicle_new**.

**Step 3: Add more proteins of a different type**

Now we also want to add some of the protein of type 2 by modifying the **point_vesicle_new**. Execute the following command:

```
TS2CG INU -p point_vesicle_new -n 2 -t 2 -r 5 -o point_vesicle_new2 -l outer
```

**NOTE:** Different values of the **–-radius** flag will alter the number of available positions to place proteins, a larger radius will result in fewer protein positions.

This command will modify **point_vesicle_new** by adding two more proteins of type 2 in the outer leaflet and create an updated folder called **point_vesicle_new2** containing the total amount of proteins. In total, the system now contains five proteins of type 1 and three proteins of type 2. The number of proteins and corresponding type can be seen in **point_vesicle_new2/IncData.dat**.

```
< Inclusion NoInc 8 >
< id typeid pointid lx ly lz >
          0            1            5    0.499   -0.864    0.059
          1            1           22    0.481   -0.765   -0.429
          2            2           30   -0.177   -0.945    0.275
          3            1         7030    1.000    0.000    0.000
          4            1          709    1.000    0.000    0.000
          5            1          241    1.000    0.000    0.000
          6            2         5382    1.000    0.000    0.000
          7            2         1555    1.000    0.000    0.000
```

**Step 4: Build a bilayer vesicle with proteins based on the modified point folder using **PCG****

Finally, in this step you will use **PCG** as done in previous tutorials but with **point_vesicle_new2** provided by **INU** in the previous step.

```
TS2CG PCG -str input_vesicle.str -Bondlength 0.2 -LLIB "./files/Martini3.LIB" -dts ./point_ves
```

The system can now be visualized using VMD or similar software, where the eight proteins can be seen as part of the membrane.

A script named **run_tut8_Vesicle.sh** is available in the **tut8/outputs/part1** folder. This script will execute the commands presented above and it will run the TS2CG output using GROMACS.

1. **Energy Minimization without Solvent**: Conduct a standard energy minimization, excluding solvent from the system.
2. **Short Equilibration without Solvent**: Run a brief equilibration step without solvent.

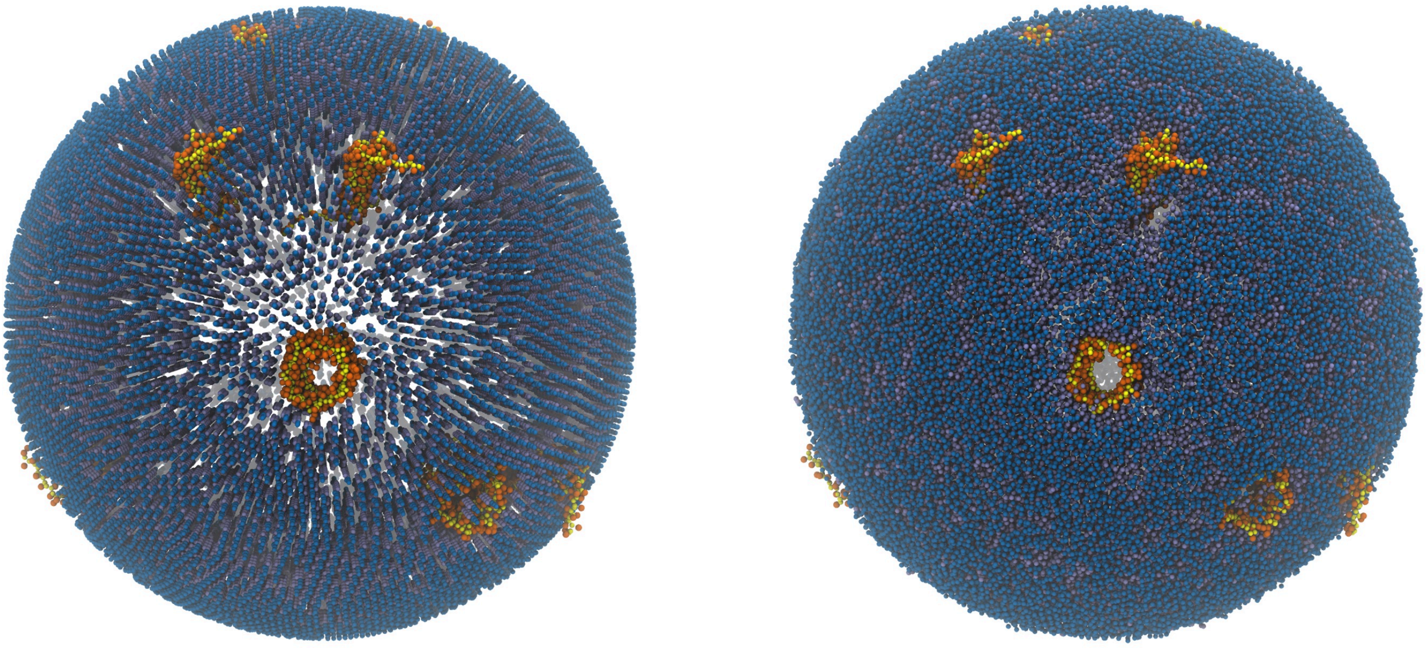

**Fig. 8-1.** Initial structure of the created POPC vesicle with eight proteins (left), and the vesicle after a brief vacuum simulation using GROMACS (right). Visualized with VMD.

**8-2: Place proteins based on curvature**

**Step 1: Generate point folder**

As in part I you start by generating a point folder but this time of a curved membrane. Execute the following command:

```
TS2CG PLM -TSfile prep/Small_Curved.tsi -bilayerThickness 3.8 -rescalefactor 4 4 4
```

**Step 2: Add proteins based on curvature**

In this tutorial, you will place the proteins in the negatively curved region by using the following command. You are placing 10 proteins of type 1 where the curvature is negative and we amplify the placement by setting k to a large number namely, 100, and the proteins will be placed in both leaflets.

Execute **INU** by using the following command:

```
TS2CG INU -p point_SmallCurved -n 10 -t 1 -r 5 -c -0.5 -k 100 -o point_SmallCurved_new -l both
```

The modifications will be saved in a new point folder called **point_new**.

**Step 3: Build a curved bilayer membrane based on the modified point folder using PCG**

**Step 4: Build a bilayer based on the modified point folder using PCG**

Finally, build the bilayer by executing the following command:

```
TS2CG PCG -str input_SmallCurved.str -Bondlength 0.2 -LLIB ./files/Martini3.LIB -dts ./point_S
```

You can now visualize the system using VMD or similar software, where you see a curved membrane with proteins in or close to the negatively curved region.

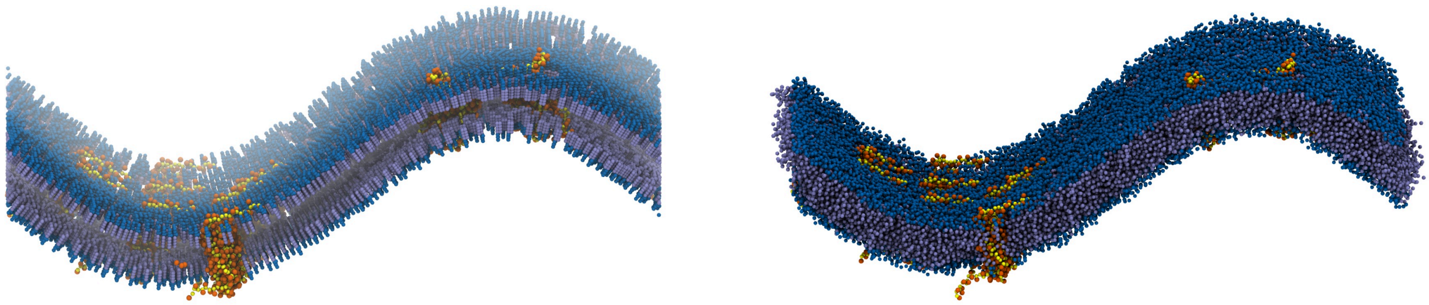

**Fig. 8-2.** Initial structure of the bilayer membrane with proteins positioned in negatively curved regions (left), and the system after a brief vacuum simulation using GROMACS (right). Visualized with VMD.

A script named **run_tut8_SmallCurved.sh** is available in the **tut8/outputs/part2** folder. This script will execute the commands presented above and it will run the TS2CG output using GROMACS.

**1. Energy Minimization without Solvent:** Conduct a standard energy minimization, excluding solvent from the system.

**2. Short Equilibration without Solvent:** Run a brief equilibration step without solvent.

#### Tutorial 9: Membrane Modifications With the Point Class [Jupyter]

The files needed for the tutorial can be manually downloaded here.

This is the notebook for Tutorial 9 of TS2CG, each python and **PCG** cell can be directly moved to a Jupiter notebook to test the steps. In this tutorial, we will use the base features of the point class in Python, which modifies the point folder created by **PLM**. After each modification, we can update the **point** folder and run **PCG** to obtain a structure.

We will read in a membrane, explore the different methods to access different parts of the folder, and modify some inclusions, exclusions, and domains.

```
# We do some imports to help us with our different steps below:

import numpy as np

# And we load TS2CG to get access to the framework

import TS2CG
```

For this first part of the tutorial, we will start with the **point** folder created in Tutorial 4. The folder contains a round bilayer vesicle and three protein inclusions of two different types.

```
point=TS2CG.core.point.Point("./point_tut4_start")
```

We are given a warning here. Luckily, this is nothing to worry about, as it just informs us that one of our input files is empty. Something we can easily check and confirm if we want it that way. In our case, **point_tutorial4/ExcData.dat** does not contain any exclusions, which is acceptable. It does, however, contain Inclusions. In this next step, we will add some more.

```
# First, let's look at the inclusions we got.

point.inclusions.get_all()

# We see a dictionary of all the inclusions, their id, their type, and their location on the p
```

```
[{'id': 0,
  'type_id': 1,
  'point_id': 5,
  'orientation': array([ 0.499, -0.864,  0.059])},
 {'id': 1,
  'type_id': 1,
  'point_id': 22,
  'orientation': array([ 0.481, -0.765, -0.429])},
 {'id': 2,
  'type_id': 2,
  'point_id': 30,
  'orientation': array([-0.177, -0.945,  0.275])}]
```

```
# Now we add another protein of type 2.

point.inclusions.add_protein(type_id=2,point_id=17)
```

```
# We can also set the orientation, but now it will default to [1,0,0].
# Let's look at our inclusions list, but restrain ourselves to proteins of type 2.

point.inclusions.get_by_type(2)
```

```
[{'id': 2,
  'type_id': 2,
  'point_id': 30,
  'orientation': array([-0.177, -0.945,  0.275])},
 {'id': 3, 'type_id': 2, 'point_id': 17, 'orientation': array([1., 0., 0.])}]
```

```
# Now we overwrite the point folder with our altered version.

point.save()

#Let's run PCG and look at the new inclusion.

!TS2CG PCG -str input.str -Bondlength 0.2 -LLIB Martini3.LIB -defout system -dts point_tut4_st
```

As we can see, we have added another protein of type 2 to our vesicle.

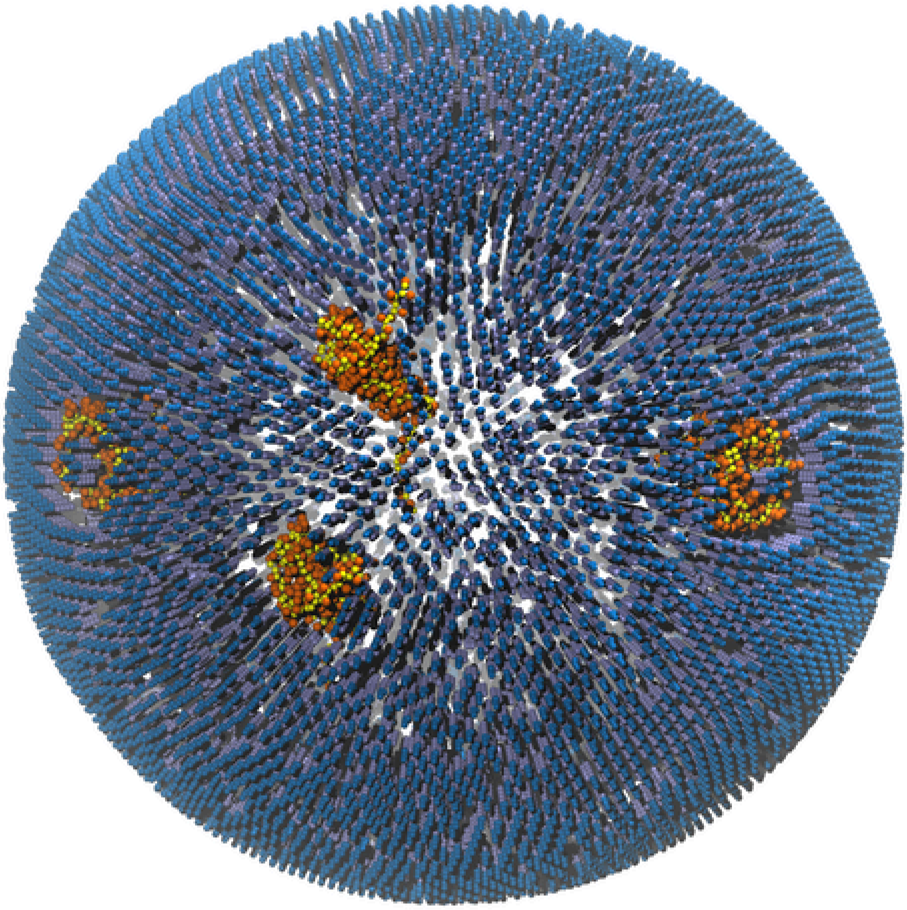

Fig. 9-1.

But, similarly to tutorial 4, lipids are in the transport protein, which was fixed via exclusions in the earlier tutorial. Let's add exclusions here as well.

```
# Let's look at the existing exclusions.

point.exclusions.get_all()

# No exclusions found. Let's loop through our proteins of type 2 and add exclusions in the exa

exclusion_locations=[d["point_id"] for d in point.inclusions.get_by_type(2)]
for id in exclusion_locations:
    point.exclusions.add_pore(point_id=id,radius=1.0)

#See, if it populated the exclusions, and save the folder a long the way for visualization

point.exclusions.get_all()
point.save()
```

```
point.exclusions.get_all()
```

```
[{'id': 0, 'point_id': 30, 'radius': 1.0},
 {'id': 1, 'point_id': 17, 'radius': 1.0}]
```

```
!TS2CG PCG -str input.str -Bondlength 0.2 -LLIB Martini3.LIB -defout system -dts point_tut4_st
```

We have successfully removed the lipids from inside the protein by overlaying the exclusions.

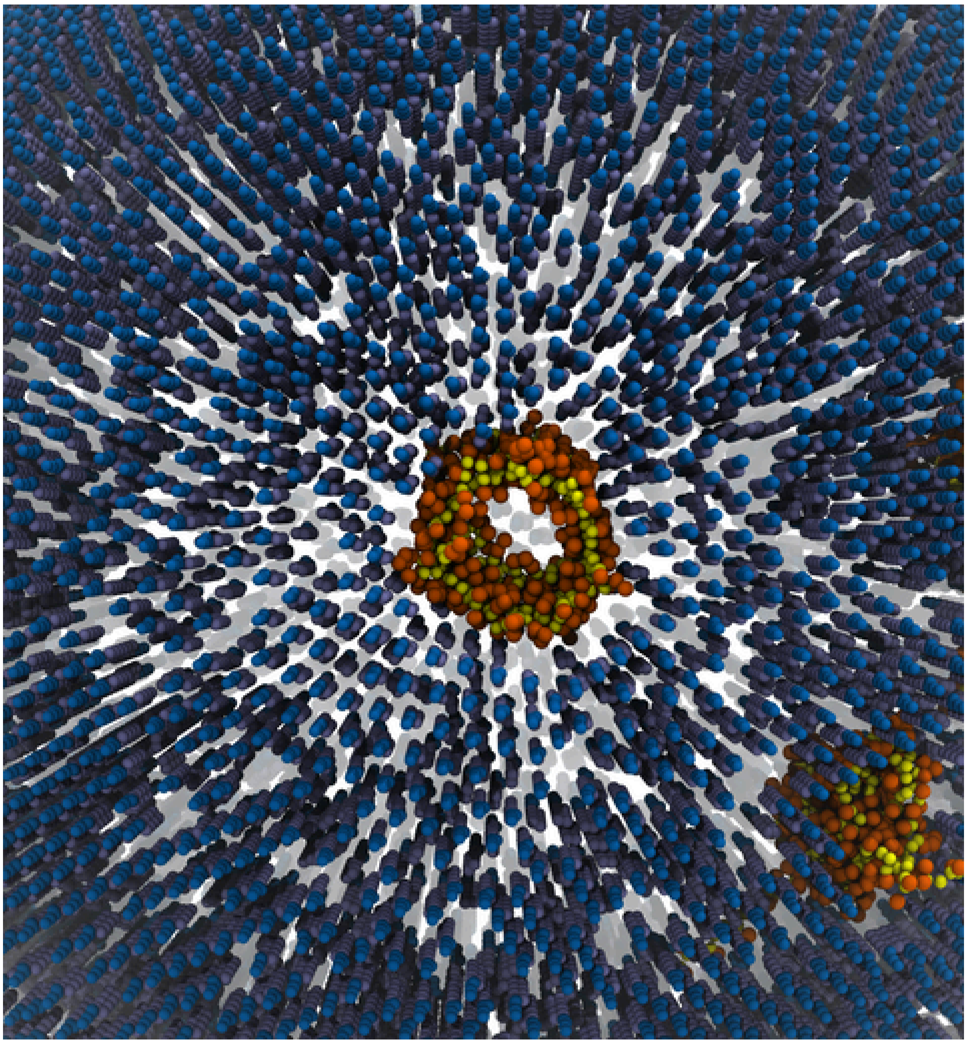
Fig. 9-2.

Now, let's add a secondary domain, somewhere.

We will want to add some POPE lipids in a weighted randomized fashion. While this might not be particularly physical, it will serve the tutorial. We will consider each point and might change the domain to 1, the standard domain is 0. For each point, we will consider the distance to the north pole. The closer to the north pole, the more likely a different domain will be assigned.

```
# We start by looping over the point and finding the north pole. The north pole will be where

id_with_maximum_z=np.argmax(point.outer.coordinates[:,2])
id_with_minimum_z=np.argmin(point.outer.coordinates[:,2])
maximum_distance=np.linalg.norm(point.outer.coordinates[id_with_maximum_z]-point.outer.coordin

for i,Entry in enumerate(point.outer.coordinates):
    distance=np.linalg.norm(Entry-point.outer.coordinates[id_with_maximum_z])
    probability = max(0, .8 - distance / maximum_distance) #The maximum chance of a POPE domai
    if np.random.rand()<probability:
        point.outer.domain_ids[i]=1

point.save()
```

In order to receive a output from PCG, we need to update the input.str and tell PCG about the new lipid we want in domain 1. This has been done in input_POPE.str So we can run PCG by just refering to the altered str.

```
!TS2CG PCG -str input_POPE.str -Bondlength 0.2 -LLIB Martini3.LIB -defout system -dts point_tu
```

This places POPE lipids accordingly.

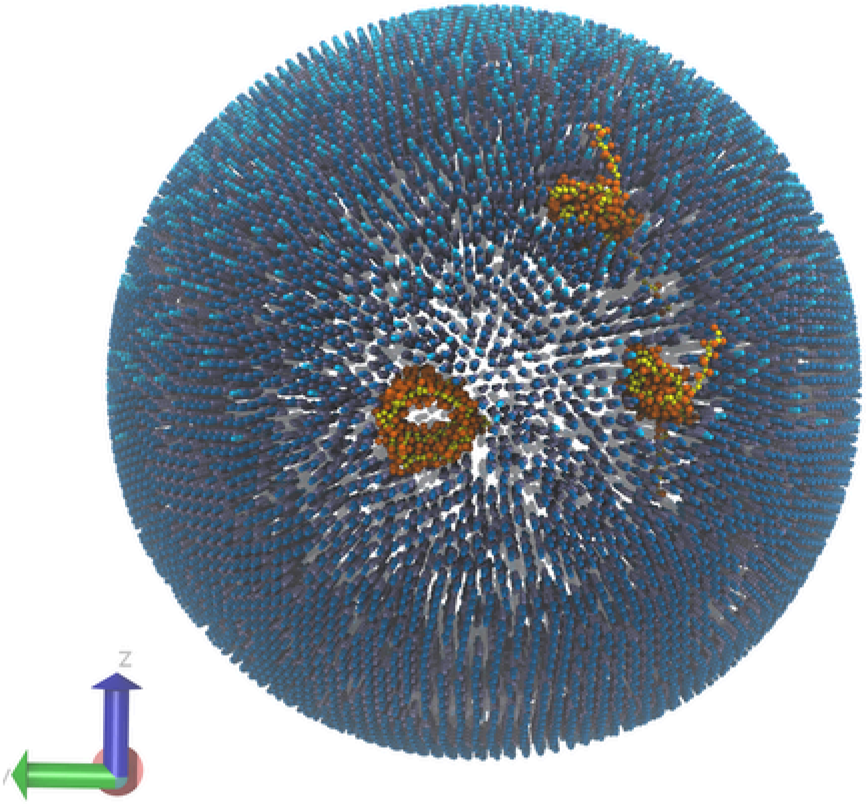


Fig. 9-3.

With this, we have used the point folder manipulator class in TS2CG to change the inclusions, the exclusions, and the domain composition on the fly and tailored to our needs.

#### Tutorial 10: Simulating a Membrane

The files needed for the tutorial can be manually downloaded here.

In this tutorial, we will solvate the 1D Fourier-shaped POPC bilayer obtained from Tutorial 6 after vacuum equilibration using the **SOL** tool and then run the solvated system with GROMACS. For this, you will need the **eq_v.gro** and **system.top** files from Tutorial 6.

Note: Before solvation, always verify that the box size matches your requirements. To perform solvation, execute the following command:

```
TS2CG SOL -in eq_v.gro -tem ./files/water.gro -o SOL.gro -Rcutoff 0.32
```

This command generates two output files:

- **SOL.gro**: The solvated structure file.
- **info.text**: Details the number of water beads to add to the topology file. After this step, the system is ready for further simulation or analysis.

A script named **run_tut10_1.sh** is provided to automate the solvation of the 1D Fourier-shaped POPC membrane and the subsequent simulations using GROMACS. The script follows these steps:

1. Standard energy minimization
2. Equilibration (with lipid head groups restrained)
3. Production run

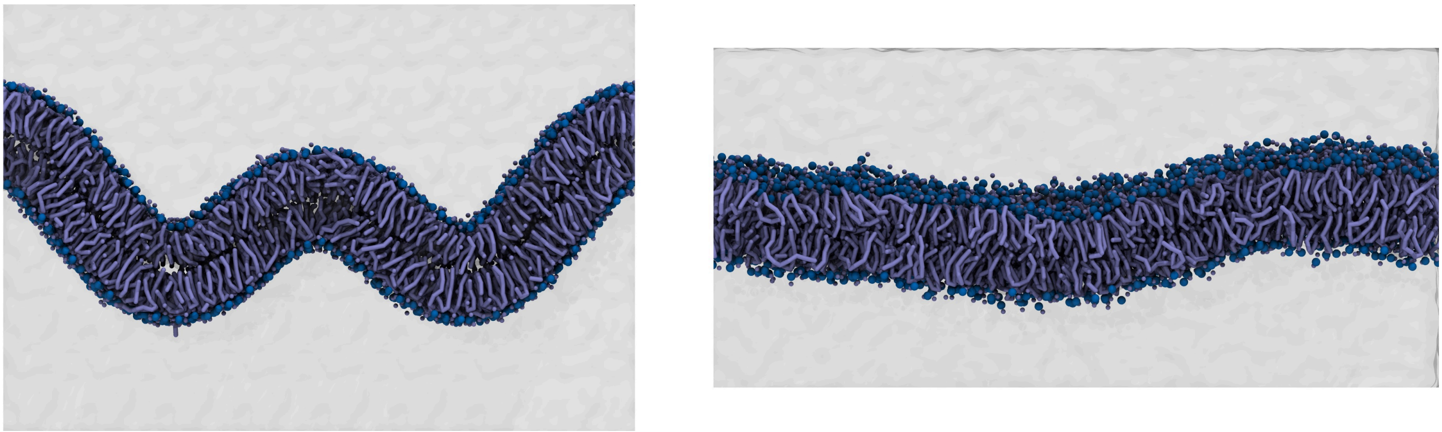

**Fig. 10-1.** Solvated 1D Fourier-shaped POPC membrane using TS2CG (left), and after simulation using GROMACS (right). Visualized with VMD.

Upon completing the production run, you will observe that the system transitions into a flat bilayer, as expected. This is where introducing a "Wall," as discussed in the second part of Tutorial 6, becomes useful.

To proceed, use **SOL** again to solvate the equilibrated (in vacuum) 1D Fourier-shaped POPC bilayer with a wall. You will need the **eq_v.gro**, **system_2.top**, and **Wall.itp** files from the second part of Tutorial 6. After solvation, simulate the system using GROMACS, following the same steps as above. The wall beads should be restrained for the energy minimization and equilibration steps.

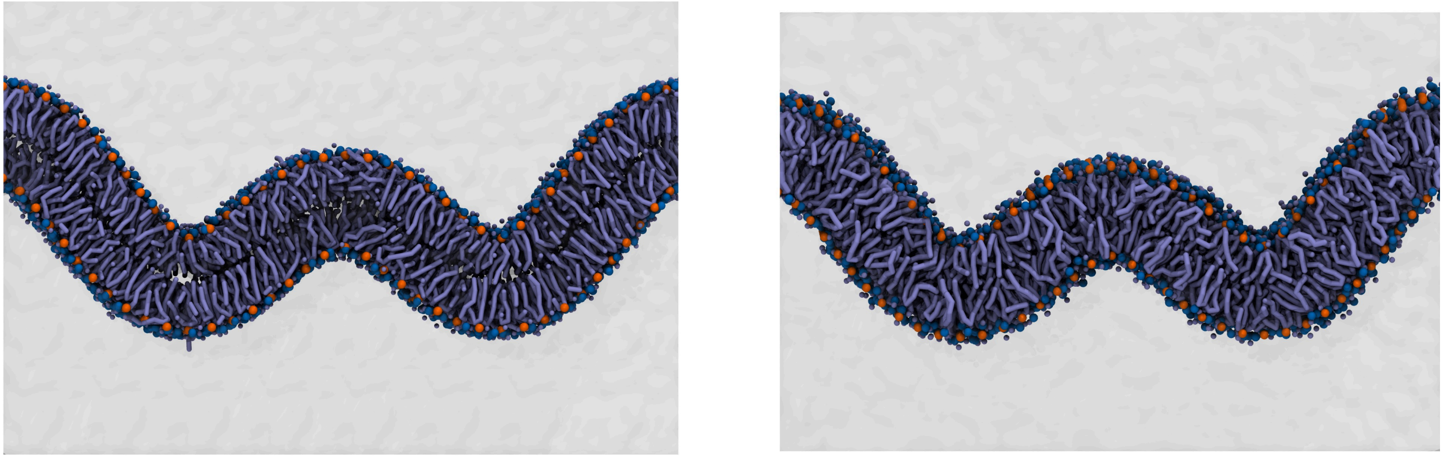

**Fig. 10-2.** Solvated 1D Fourier-shaped POPC membrane with a wall using TS2CG (left), and after simulation using GROMACS (right). Visualized with VMD.
