## Supplementary material for "TS2CG as a membrane builder": Details to S2

### TS2CG version 2.0

TS2CG converts triangulated surfaces (TS) to coarse-grained membrane models for molecular simulation. It also works as a backmapping algorithm from dynamically triangulated surfaces simulations to CG molecular dynamics simulations or to take electron microscopy tomography data and build structures for molecular dynamics simulations.

#### State

**NOTE**: this version is under development and might be associated with errors and bugs. Please use the previous version if you are not in direct contact with the developers. Previous version can be found here.

#### Installation

##### Prerequisites

TS2CG is implemented in C++ and includes three separate scripts. *Pointillism* (PLM) and *Membrane Builder* (PCG) and *Solvate* (SOL).

The minimum installation requirements are:

- Up-to-date *C++ compilers*.
- *Python* 3.6 or later.
- *CMake* version 3.10 or later.

TS2CG builds with the CMake build system, requiring at least version 3.10. You can check whether CMake is installed, and what version it is, with

```
cmake --version
```

If you need to install CMake, then first check whether your platform's package management system provides a suitable version, or visit the CMake installation page for pre-compiled binaries, source code and installation instructions.

##### Install *TS2CG*

###### From PyPi

```
pip3 install TS2CG
```

###### Directly from GitHub

```
pip3 install git+https://github.com/weria-pezeshkian/TS2CG-v2.0
```

###### From source

```
git clone https://github.com/weria-pezeshkian/TS2CG-v2.0
cd TS2CG-v2.0
python3 -m venv venv && source venv/bin/activate # Not required, but often convenient.
pip3 install .
```

#### Usage

For help on TS2CG and it's executables run:

```
TS2CG -h
TS2CG {SOL,PLM,PCG} -h
```

### Pointillism (PLM)

Pointillism (PLM), reads a triangulated surface input file e.g., typical outputs of dynamically triangulated surfaces simulations, and generates two sets of points that represent a two smooth surfaces (upper and lower monolayers of a bilayer). It extends the TS file based on the given rescaling factor and an approximated area per lipid.

##### Input and output files

The PLM executable requires a triangulated surface file as input.

A triangulated surface file that can be read by this script should be one of the following files:

- `.tsi` (see .tsi file)

- `.q` (see .q file)
- `.dat` (see .dat file)

Output files will be separated into two folders.

- A folder containing a few visualization files.
- A folder that can be read by CG Membrane Builder script (see below).

#### Command line options

| Option | Type | Default | Description |
|---|---|---|---|
| `-rescalefactor` | rx ry rz | (1 1 1) | Rescaling factor |
| `-bilayerThickness` | double | 3.8 | Bilayer thickness |
| `-monolayer` | int | 0 | To generate monolayer instead (1/-1). |
| `-r` | string | PLM | Function (PLM/in_out/check/add_pbc). |
| `-smooth` | ------ | no | Might be necessary for rough surfaces. |
| `-o` | string | point | Name of the output folder. |
| `-resizebox` | ------ | no | Find a better box for the system. |
| `-TSfile` | string | TS.tsi | TS file name (three file format types: *.q, *.tsi, *.dat). |
| `-Mashno` | int | 1 | Number of Mosaicing, your point number grows as 4^Mashno. |
| `-AlgType` | string | Type1 | Algorithm type for Mosaicing (Type1 and Type2); no difference has been reported yet. |

#### Notes

- The approximated area per lipid does not need to be precise, it will be modified during the later processes.
- The number of the output points is always larger or equal the number of the vertices in the input triangulated surface. With option `-Mashno` you can tune how many points you want (No_of_vertex*4^Mashno).
- There is no guarantee to get proper surface if rx, ry, rz are not equal.
- Use Mashno [1-4], unless you know what you are doing.

#### Usage example

```
TS2CG PLM -TSfile Traj1.tsi -bilayerThickness 4 -rescalefactor 3 3 3 -o output
```

### Membrane Builder (PCG)

Membrane builder generates a Gromacs based topology and coordinate file using the Pointilism output files. If the system contains protein, or if the user wants to add proteins, a Gromacs coordinate file (.gro file) of the protein structure needs be provided. By default, Martini forcefield lipids will be used. Read more about the Martini forcefield on the Martini website. To backmap to a CG forcefield a lipid structure library needs be provided.

#### Input and output files

The PCG executable requires an input file with an .str extension (see input.str file).

There are 3 output files generated by PCG:

- A pcg.log file containing information about the PCG execution.
- A .gro file containing the coordinates of the generated system (see gro file).
- A .top file describing the composition of the generated system (see top file).

#### Command line options

| Option | TypAe | Default | Description |
|---|---|---|---|
| `-dts` | string | point | DTS folder that was generated by PLM |
| `-str` | string | input.str | Input file |
| `-defout` | string | output | Output files prefix |
| `-Bondlength` | double | 0.1 | Initial bond guess |
| `-LLIB` | string | Martini3 | CG lipid library file name |
| `-renorm` | ------ | no | Renormalizes the lipid molar ratio |
| `-iter` | double | 4 | The number of point selections is `iter * number of the point` |
| `-incdirtype` | string | Global | The type of protein direction data ( `Local` or `Global` ) |
| `-Wall` | ------ | off | A flag to create a wall around the membrane |
| `-function` | string | backmap | Backmap or analytical shape |

| Option | TypAe | Default | Description |
|---|---|---|---|
| `-WallBName` | string | WL | Name of the Wall beads |
| `-WPointDir` | bool | false | Just write the folder |

#### Notes

- With option `-Bondlength`, you can change the initial bond guess. Large Bondlength may generate an unstable structure.
- With option `-renorm` the molar ratio of the lipid will be renormalized.
- To get higher denisty, you may increase `-Mashno` value or reduce -ap value in PLM command.
- When using `-function analytical shape`, the input.str file must contain the [Shape Data] section (see input.str file).

#### Usage example

```
TS2CG PCG -dts point -str input.str -seed 39234  -Bondlength 0.15
```

### Python Documentation

The Point class is the core of TS2CG regarding the Python scripts. The documentation of these python modules and the point class can be found here: TS2CG 2.0 - Python Documentation
You can find the python documentation below on this document.

### Solvation (SOL)

The Solvation executable solvates the system and provides an option to add ions.

For information on the script run:

```
TS2CG SOL -h
```

#### Input and output files

The SOL executable requires the input file of the solvent with a .gro extension (see gro file). The default option is `input.gro`.

There are 3 output files generated by SOL:

- An `info.txt` file containing the amount of atoms added to the system. The content of the info.txt file should be appended to the topology file. Also remember to include the necessary itp

files for solvents and/or ions.
- A `.gro` file containing the coordinates of the solvated system (see gro file).

##### Command line options

| Option | Type | Default | Description |
|---|---|---|---|
| `-in` | string | `input.gro` | Name of the file to be solvated |
| `-ion` | integers | `0 0` | Generating ions; two integers: number of positive and negative ions |
| `-db` | double | `0.05` | The distance between each copy when propagating the water box |
| `-o` | string | `output.gro` | Output file name |
| `-Rcutoff` | double | `0.4` | Cutoff distance |
| `-tem` | string | `W.gro` | Name of the template file for solvation |
| `-nname` | string | `CL` | Name of the negative ions |
| `-pname` | string | `NA` | Name of the positive ions |
| `-seed` | integer | `9474` | Seed for ion placement |
| `-unsize` | double | `2` | Size of the unit cells for overlap checking; smaller numbers are faster but need more RAM. Should not be smaller than `Rcutoff`. |

##### Usage example

```
SOL -in in.gro -o out.gro -ion 20 20 -tem water.gro
```

### File formats

#### q file

The q file can be used as input to the PLM executable and is formatted as shown below:

```
50.000      50.000      50.000
1840
0        21.4     33.8     32.7     0
1        38.1     26.1     32.3     0
```

```
2       40.9    24.2    19.9    0
...
1839    31.2    323.2   23      0
3680
0        75      776    1043    1
1       796     1821     752    1
2       995     1027     279    1
3       662    1162      56     1
4       167      38     391     1
...
```

- Line 1: Box information (3 double numbers).
- Line 2: Number of vertices (1 integer number; Let's call it NV).
- Line 3 to NV+2: Vertex ID, coordinates (1 integer number and 3 double numbers).
- Line NV+3: Number of triangles (1 integer number; Let's call it NT).
- Line NV+4 to NV+NT+3: Triangle ID, ID of its vertices, and type (5 integer numbers).

##### Notes

- The last integer in each row of vertex and triangles is the type of the specific object. - This type of input file does not support inclusions.

#### tsi file

The `*.tsi` files, are DTS simulation trajectory outputs. It contains information about vertices, triangles and inclusion positions.
It can be used as input to the PLM executable and it is formated as shown below:

###### General Structure

1. Each `.tsi` file begins with a line specifying **version 1.1**.
2. The next line defines the **box size** (`x`, `y`, and `z`) of the system in **nm**.
3. The subsequent three sections describe the **TS mesh**. Each section starts with a keyword (`vertex`, `triangle`, or `inclusion`) and their respective counts.

```
version 1.1
box 50.000     50.000     50.000
vertex 1840
0       21.4    33.8    32.7    0
1       38.1    26.1    32.3    0
2       40.9    24.2    19.9    0
...
1839    31.2    323.2   23      0
triangle 3680
0        75      776    1043    1
1       796     1821     752    1
2       995     1027     279    1
```

```
3       662    1162      56     1
4       167      38     391     1
...
inclusion 3
0         1       22       0    1
1         1        5       0    1
2         2       30       0    1
```

##### Vertex section

- The file includes **1840 vertices**.
- Each vertex is assigned:
  - An **index**.
  - A **position** in `x`, `y`, and `z`.
  - An integer representing the **domain**.

##### Triangles

- The 1840 vertices are connected via **3680 triangles**.
- Each triangle is defined by:
  - An **index**.
  - The **vertices** it connects.
  - An integer representing the **type**.
- Example:
  - Triangle `0` connects vertices `75`, `776`, and `1043`.

##### Inclusions

- A `.tsi` file may include a section for **(protein) inclusions**.
- In this example:
  - There are **three inclusions** of **two different types**.
- Each inclusion is defined by:
  i. An **index**.
  ii. The **inclusion type** (e.g., type `1` for inclusions `0` and `1`, type `2` for inclusion `2`).
  iii. The **corresponding vertex index**.
  iv. **Two floating-point numbers**:
    - These describe a unit two-dimensional vector.
    - The numbers sum to **1**.
    - They define the **orientation** of the inclusion relative to the bilayer normal.

### gro file

The structure of the gro file can be seen below:

```
TS2CG manual
    6
    1WATER  OW1    1   0.126   1.624   1.679  0.1227 -0.0580  0.0434
    1WATER  HW2    2   0.190   1.661   1.747  0.8085  0.3191 -0.7791
    1WATER  HW3    3   0.177   1.568   1.613 -0.9045 -2.6469  1.3180
    2WATER  OW1    4   1.275   0.053   0.622  0.2519  0.3140 -0.1734
    2WATER  HW2    5   1.337   0.002   0.680 -1.0641 -1.1349  0.0257
    2WATER  HW3    6   1.326   0.120   0.568  1.9427 -0.8216 -0.0244
   1.82060   1.82060   1.82060
```

- Line 1: Title (free format string).
- Line 2: Number of atoms (1 integer number; Let's call it NA).
- Line 3 to NA+2: Each column corresponds to: Residue Number | Residue Name | Atom Name | Atom Number | Positions (x, y, z) | Velocities (vx, vy, vz).
- Line NA+3: Box vectors (free format, space-separated reals).

For more information on the gro file format and usage see Gromacs Documentation.

### input.str file

Files with str extension are used as input files in the PCG executable and are structured as below:

```
include protein.gro
[Lipids List]
;LipidName  RatioUp RatioDown   Area/Lipid
Domain 0
POPC        0.5     0.5         0.63
POPE        0.5     0.5         0.64
End
Domain 1
POPC        0.5     0.5         0.63
POPE        0.5     0.5         0.64
End
[Shape Data]
ShapeType     Flat
Box    40     40     40
Density       2      2
Thickness     3.8
WallRange     0      1    0    1
End
[Protein List]
;ProteinName    Incl.Id     Surface_Coverage    0   0   z-position
STxB            1           0.5                 0   0   -1.1
end Protein
```

- Inclusions of necessary files. One line for each file.
- Initialization of [Lipid list] section.
- Initialization of Domain.
- Lipid lines: Lipid Name | Ratio Up | Ratio Down | Area per lipid in nm-2. (1 string and 3 float numbers).
- Termination of Domain.
- Initialization of [shape data] section. This section should exist when using the flag `-function analytical_shape`. You can see the rest of the analytical shapes in Tutorial 6
- Initialization of [Protein list] section.
- Protein lines: ProteinName | Inclusion ID | Surface Coverage | 0 | 0 | z-position.
- Termination of Protein list.
- The comments in the **input.str** file is initiated with a semicolon ";".

#### top file

```
;
;       Example topology file
;
; The force-field files to be included
#include "amber99.ff/forcefield.itp"
[ moleculetype ]
; name  nrexcl
Urea         3
[ atoms ]
   1  C  1  URE      C      1     0.880229  12.01000   ; amber C  type
   2  O  1  URE      O      2    -0.613359  16.00000   ; amber O  type
   3  N  1  URE     N1      3    -0.923545  14.01000   ; amber N  type
   4  H  1  URE    H11      4     0.395055   1.00800   ; amber H  type
   5  H  1  URE    H12      5     0.395055   1.00800   ; amber H  type
   6  N  1  URE     N2      6    -0.923545  14.01000   ; amber N  type
   7  H  1  URE    H21      7     0.395055   1.00800   ; amber H  type
   8  H  1  URE    H22      8     0.395055   1.00800   ; amber H  type
[ bonds ]
    1    2
    1    3
    1    6
    3    4
    3    5
    6    7
    6    8
[ dihedrals ]
;   ai     aj     ak     al  funct   definition
     2      1      3      4    9
     2      1      3      5    9
     2      1      6      7    9
     2      1      6      8    9
     3      1      6      7    9
```

```
     3     1     6     8   9
     6     1     3     4   9
     6     1     3     5   9
[ dihedrals ]
     3     6     1     2   4
     1     4     3     5   4
     1     7     6     8   4
[ position_restraints ]
; you wouldn't normally use this for a molecule like Urea,
; but we include it here for didactic purposes
; ai   funct    fc
   1     1     1000    1000    1000 ; Restrain to a point
   2     1     1000       0    1000 ; Restrain to a line (Y-axis)
   3     1     1000       0       0 ; Restrain to a plane (Y-Z-plane)
[ dihedral_restraints ]
; ai   aj    ak    al  type  phi  dphi  fc
    3    6     1    2     1  180     0  10
    1    4     3    5     1  180     0  10
; Include TIP3P water topology
#include "amber99.ff/tip3p.itp"
[ system ]
Urea in Water
[ molecules ]
;molecule name   nr.
Urea             1
SOL              1000
```

- #include “amber99.ff/forcefield.itp” : this includes the information for the force field you are using, including bonded and non-bonded parameters.
- [ moleculetype ] : defines the name of your molecule in this top and nrexcl = 3 stands for excluding non-bonded interactions between atoms that are no further than 3 bonds away.
- [ atoms ] : defines the molecule, where nr and type are fixed, the rest is user defined. So atom can be named as you like, cgnr made larger or smaller (if possible, the total charge of a charge group should be zero), and charges can be changed here too.
- [ bonds ] : The pairs of bonds between the atoms of the molecule.
- [ dihedrals ] : in this case there are 9 proper dihedrals (funct = 1), 3 improper (funct = 4) and no Ryckaert-Bellemans type dihedrals.
- [ position_restraints ] : harmonically restrain the selected particles to reference positions (Position restraints).
- [ dihedral_restraints ] : restrain selected dihedrals to a reference value.
- #include “tip3p.itp” : includes a topology file that was already constructed
- [ system ] : title of your system, user-defined
- [ molecules ] : this defines the total number of (sub)molecules in your system that are defined in this top.

The line of a comment on a topology file should start with with a semicolon ";" .
For more deltailed explaination of the structure and usage of a **.top** file see Gromacs Documentation.

#### LIB file

Lipid library files with `.LIB` extension are used as input in the PCG executable and are structured as below:

```
Description Martini Map CG
Version Martini 3
;Martini 3 lipid library for TS2CG 2.0
;Made by Reinier de Vries on 31-03-2021. Last updated: 04-05-2021
;NOTE: Bead atom names don't match current Martini 3 bead atom names, use -maxwarn in GROMACS
to ignore this. (Inconsistency in naming of tail beads in Martini 3)
[DAPC]
1 NC3 0 0    1
2 PO4 0 0    0
3 GL1 0 0   -1
4 GL2 0 0.5 -1
5 D1A 0 0 -2
6 D2A 0 0 -3
7 D3A 0 0 -4
8 D4A 0 0 -5
9 C5A 0 0 -6
10 D1B 1 0 -2
11 D2B 1 0 -3
12 D3B 1 0 -4
13 D4B 1 0 -5
14 C5B 1 0 -6


[POPC]
1 NC3 0 0    1
2 PO4 0 0    0
3 GL1 0 0   -1
4 GL2 0 0.5 -1
5 C1A   0     0   -2
6 D2A   0     0   -3
7 C3A   0     0   -4
8 C4A   0     0   -5
9 C1B   1     0   -2
10 C2B  1     0   -3
11 C3B  1     0   -4
12 C4B  1     0   -5
```

- Each section on the file represents a lipid, starting by the name in square brackets.

- The rest fo the lines in a section represent each individual bead. The lines are as follows: Bead Line: Internal ID | Name | x-offset | y-offset | z-offset.
- Theese offsets are then multiplied with the BondLenght to place the beads onto the correct relative position.

The line of a comment on a topology file should start with with a semicolon ";" .
You can find a Martini3.LIB file with the most common lipids on the files of the tutorials (.LIB file).

#### dat files

#### Quick references

About TS2CG
New Updates
Tutorials for version 2.0
Tutorials 1.1
