## Supplementary material for "TS2CG as a membrane builder": Details to S3

### TS2CG Version 2 - Python documentation

Welcome to the python documentation of the second version of TS2CG. Here, you can find details about the *Core* classes and their functionality as well as information on the usage of the python tools included in TS2CG. NOTE: This version might be associated with errors and bugs. Please use the previous version if you are not in direct contact with the developers. Previous version can be found here: marrink-lab/TS2CG1.1 For documentation on the rest of the modules of TS2CG see: weria-pezeshkian/TS2CG-v2.0

This documentation is divided into two sections:

- **Core**: Details on the functionality of the *Core* classes.
- **Tools**: Explains the individual tools (*DAI*, *DOP*, *INU*, *VIS*).

Contents:


- Core Documentation
  - Exclusion class
  - Inclusion class
  - Membrane class
  - Point class
- Tools Documentation
  - **DAI**
  - **DOP**
  - **INU**
  - **VIS**


Next
Core Documentation

Exclusion class

Inclusion class

Membrane class

Point class

### Core Documentation

Contents:

- Exclusion class
- Inclusion class
- Membrane class
- Point class

Previous
TS2CG Version 2 - Python documentation

Next
Exclusion class

Exclusion class
Inclusion class
Membrane class
Point class

### Exclusion class

`class TS2CG.core.exclusion.Exclusion(data: ndarray | None = None)` [source]

Bases: `object`

Manages lipid exclusions in the membrane.

`add_pore(point_id: int, radius: float = 1.0)` [source]

Add a pore in the lipid membrane.

**Args:**

point_id: Point ID where lipids should be excluded radius: Radius of exclusion zone

`get_all() → List[int]` [source]

Get all excluded exclusions.

`remove_pore(index: int)` [source]

remove a pore in the lipid membrane.

**Args:**

**index: int**

index of exclusion to remove

`remove_pores(indices: List[int])` [source]

remove a list of pores from the lipid membrane.

**Args:**

**indices: List[int]**

list of indices of inclusions to remove

Search the docs ...

CONTENTS:


Exclusion class

Inclusion class

Membrane class

Point class


### Inclusion class

*class* TS2CG.core.inclusion.**Inclusion**(*data: ndarray | None = None*) [source]

Bases: `object`

Manages protein inclusions in the membrane.

**add_protein**(*type_id: int, point_id: int, orientation: ndarray | None = array([1, 0, 0])*) [source]

Add a protein inclusion.

**Args:**

type_id: Type identifier for the protein point_id: Point ID where protein should be placed orientation: Vector specifying protein orientation

**get_all**() → List[int] [source]

Get all inclusion with protein inclusions.

**get_by_type**(*type_id: int*) → List[dict] [source]

Get all inclusions of a specific type.

**remove_protein**(*index: int*) [source]

remove a protein inclusion.

**Args:**

**index: int**

index of inclusion to remove

**remove_proteins**(*indices: List[int]*) [source]

remove a list of protein inclusions.

**Args:**

**indices: List[int]**

list of indices of inclusions to remove


Previous
TS2CG.core.exclusion module

Next
Membrane class

Search the docs ...

CONTENTS:

Exclusion class

Inclusion class

Membrane class

Point class

### Membrane class

*class* TS2CG.core.membrane.**Membrane**(*data: ndarray*) [source]

Bases: `object`

Represents a membrane layer with associated properties.

*property* **gaussian_curvature**: *ndarray*

Calculate Gaussian curvature for all points.

**get_edge_ids**() → ndarray [source]

Get ids of all points that are an edge.

**get_points_by_domain**(*domain_id: int*) → ndarray [source]

Get coordinates of all points in a specific domain.

*property* **mean_curvature**: *ndarray*

Calculate mean curvature for all points.

Previous
Inclusion class

Next
Point class

Search the docs ...

CONTENTS:


Exclusion class

Inclusion class

Membrane class

Point class


### Point class

`class TS2CG.core.point.Point(path: str | Path)` [source]

Bases: `object`

A class representing a membrane structure with inclusions and exclusions. Can be initialized from a point folder or built from scratch.

`save(output_path: str | Path | None = None, backup: bool | None = True)` [source]

Save membrane structure to files.

**Args:**

**output_path: pathlib**

Path where to save the point folder. If None, saves to original location. Backup is only created if saving to the original location.

**backup: bool**

wheter or not to write output files

`update_domains(domain_ids: ndarray | None = None)` [source]

Update domain assignments for membrane layer(s). For bilayers, updates both leaflets. For monolayers, updates only the outer leaflet.

**Args:**

domain_ids: New domain assignments as numpy array

`TS2CG.core.point.loadtxt_fix(filename, skiprows)` [source]


Previous
Membrane class

Next
Tools Documentation

Search the docs ...


CONTENTS:

DAI

DOP

INU

VIS


### Tools Documentation

The *Tools* module provides the following tools:

- **DAI**: Handles circular domains.
- **DOP**: Manages domain placement.
- **INU**: Updates inclusions.
- **VIS**: Visualizes directories.

Contents:


- DAI
- DOP
- INU
- VIS



Previous
TS2CG.core.point module

Next
DAI

Search the docs ...

CONTENTS:


DAI

DOP

INU

VIS


### DAI

CLI tool to place lipids to assign circular domains around inclusions or points.

TS2CG.tools.circular_domains.DAI(*args: List[str]*) → None [source]

Main entry point for Domain Placer tool

*class* TS2CG.tools.circular_domains.LipidSpec(*domain_id: int*, *name: str*, *percentage: float*, *curvature: float*, *density: float*) [source]

Bases: object

Specification for a lipid type and its properties

curvature*: float*

density*: float*

domain_id*: int*

name*: str*

percentage*: float*

TS2CG.tools.circular_domains.circular_domains(*membrane: Point*, *radius: float*, *pointid: list*, *domain: int*, *path_dist: bool = False*, *percent: float = 100.0*, *layer: str = 'both'*) → None [source]

Assign lipids to domains based on curvature preferences

TS2CG.tools.circular_domains.parse_lipid_file(*file_path: Path*) → List[LipidSpec] [source]

Parse lipid specification file into structured data

TS2CG.tools.circular_domains.write_input_str(*lipids: Sequence[LipidSpec]*, *output_file: Path*, *old_input: Path | None = None*) → None [source]

Write input.str file for TS2CG, preserving all comments and sections except [Lipids List]. Maintains exact formatting of the original file.


Previous
Tools Documentation

Next
DOP

Search the docs ...

CONTENTS:


DAI

**DOP**

INU

VIS


### DOP

CLI tool to place lipids in membrane domains based on local curvature preferences. Uses domain_input.txt format for lipid specifications and generates input.str for next steps.

Example domain_input.txt: ; domain lipid percentage c0 density 0 POPC .5 0.179 0.64 2 POPG .5 0.629 0.64

`TS2CG.tools.domain_placer.DOP(args: List[str]) → None` [source]

Main entry point for Domain Placer tool

`class TS2CG.tools.domain_placer.LipidSpec(domain_id: int, name: str, percentage: float, curvature: float, density: float)` [source]

Bases: `object`

Specification for a lipid type and its properties

`curvature: float`

`density: float`

`domain_id: int`

`name: str`

`percentage: float`

`TS2CG.tools.domain_placer.assign_domains(membrane: Point, lipids: Sequence[LipidSpec], layer: str = 'both', k_factor: float = 1.0, area: bool = False) → None` [source]

Assign lipids to domains based on curvature preferences

`TS2CG.tools.domain_placer.calculate_curvature_weights(local_curvature: float, lipids: Sequence[LipidSpec], k_factor: float, area: float = 1.0, max_delta: float = 5.0) → ndarray` [source]

Calculate Boltzmann weights for each lipid type at given curvature

`TS2CG.tools.domain_placer.parse_lipid_file(file_path: Path) → List[LipidSpec]` [source]

Parse lipid specification file into structured data

`TS2CG.tools.domain_placer.write_input_str(lipids: Sequence[LipidSpec], output_file: Path, old_input: Path | None = None) → None` [source]

Write input.str file for TS2CG, preserving all comments and sections except [Lipids List]. Maintains exact formatting of the original file.


Previous
DAI

Next
INU

DAI
DOP
INU
VIS

### INU

CLI tool to place protein inclusions in membrane. Reads input.str for protein definitions and places new inclusions in the membrane.

**Usage:**

TS2CG INU -p point -t 0 -r 2 -N 10 -c 0.1 -o point_new -l both

`TS2CG.tools.inclusion_updater.INU(args: List[str]) → None` [source]

Main entry point for protein inclusion tool

`TS2CG.tools.inclusion_updater.calculate_curvature_weights(curvatures: ndarray, target_curvature: float | None, k_factor: float) → ndarray` [source]

Calculate Boltzmann weights based on curvature preference

`TS2CG.tools.inclusion_updater.get_nearby_points_both_leaflets(membrane: Point, leaflet: str, point_idx: int, radius: float) → Dict[str, ndarray]` [source]

Find points within radius in both leaflets from a point in the specified leaflet.

**Args:**

membrane: Membrane Point object leaflet: Which leaflet the center point is in ('inner' or 'outer') point_idx: Index of the center point radius: Exclusion radius

**Returns:**

Dict with excluded points for each leaflet

`TS2CG.tools.inclusion_updater.get_points_near_existing_proteins(membrane: Point, radius: float) → Dict[str, Set[int]]` [source]

Get points that are too close to existing proteins in any leaflet

`TS2CG.tools.inclusion_updater.pbc_wrap(membrane)` [source]

Wrap membrane coordinates into the primary box

`TS2CG.tools.inclusion_updater.place_proteins(membrane: Point, type_id: int, radius: float, num_proteins: int | None = None, target_curvature: float | None = None, k_factor: float = 1.0, leaflet: str = 'both') → Dict[str, int]` [source]

Place proteins in membrane with given constraints

### VIS

CLI tool to place lipids to assign circular domains around inclusions or points.

`TS2CG.tools.dir_visualizer.VIS(args) → None` [source]

Main entry point for Domain Placer tool

`TS2CG.tools.dir_visualizer.draw_folder(membrane: Point, pointid: list, domain: bool, layer: str = 'both', save=None, step=1, Proteins=False) → None` [source]

Assign lipids to domains based on curvature preferences

`TS2CG.tools.dir_visualizer.make_fullscreen()` [source]

Make the Matplotlib figure full screen universally across platforms.
